# Mechanism of Tripeptide Trimming by γ-Secretase

**DOI:** 10.1101/2021.08.05.455243

**Authors:** Apurba Bhattarai, Sujan Devkota, Hung Nguyen Do, Jinan Wang, Sanjay Bhattarai, Michael S. Wolfe, Yinglong Miao

## Abstract

The membrane-embedded γ-secretase complex processively cleaves within the transmembrane domain of amyloid precursor protein (APP) to produce 37-to-43-residue amyloid β-peptides (Aβ) of Alzheimer’s disease (AD). Despite its importance in pathogenesis, the mechanism of processive proteolysis by γ-secretase remains poorly understood. Here, mass spectrometry and western blotting were used to quantify the efficiency of the first tripeptide trimming step (Aβ49→Aβ46) of wildtype (WT) and familial AD (FAD) mutant APP substrate. In comparison to WT APP, the efficiency of this first trimming step was similar for the I45F, A42T and V46F APP FAD mutants, but substantially diminished for the I45T and T48P mutants. In parallel with biochemical experiments, all-atom simulations using a novel Peptide Gaussian accelerated molecular dynamics (Pep-GaMD) method were applied to investigate tripeptide trimming of Aβ49 by γ-secretase. The starting structure was active γ-secretase bound to Aβ49 and APP intracellular domain (AICD), as generated from our previous study that captured activation of γ-secretase for the initial endoproteolytic cleavage of APP (Bhattarai *et al*., *ACS Cent Sci*, 2020, 6:969-983). Pep-GaMD simulations captured remarkable structural rearrangements of both the enzyme and substrate, in which hydrogen-bonded catalytic aspartates and water became poised for tripeptide trimming of Aβ49 to Aβ46. These structural changes required a positively charged N-terminus of endoproteolytic coproduct AICD, which could dissociate during conformational rearrangements of the protease and Aβ49. The simulation findings were highly consistent with biochemical experimental data. Taken together, our complementary biochemical experiments and Pep-GaMD simulations have enabled elucidation of the mechanism of tripeptide trimming by γ-secretase.

**Significance statement:** Production of amyloid β-peptide (Aβ) of Alzheimer’s disease (AD) from its precursor protein requires a series of proteolytic events carried out by the membrane-embedded γ-secretase complex. Mutations in the substrate and enzyme that produce Aβ cause hereditary AD and these mutations affect tripeptide trimming of initially formed long Aβ peptides by γ-secretase. Little is known about the structural mechanism of this trimming process. Here, we have developed a molecular dynamic model for the first step of trimming (Aβ49 to Aβ46) by γ-secretase. Conformational changes in the enzyme-substrate complex to set up this trimming step require the presence of N-terminally charged endoproteolytic cleavage co-product. Computational effects of AD-causing mutations in Aβ49 on the trimming process were validated through biochemical experiments.

## Introduction

Alzheimer’s disease (AD) contributes to more than 80% of all dementia cases (1). Deaths related to AD in the United States increased by 89% between 2000 and 2014, and more than 6.2 million Americans are affected with AD in 2021 (www.alz.org). AD is characterized by deposition of longer amyloid β-peptides (Aβ) in the form of cerebral plaques. The amyloid β-protein precursor (APP) is successively processed by β-secretase and γ-secretase to produce Aβ peptides. β-Secretase first sheds the APP extracellular domain to produce transmembrane peptide C99, followed by processive proteolysis by γ-secretase to produce Aβ peptides of varying lengths (2). Membrane-embedded γ-secretase is a multi-domain aspartyl protease with presenilin as the catalytic subunit. γ-Secretase is considered “the proteasome of the membrane”, with more than 100 known substrates, including APP and the Notch family of cell-surface receptors (3, 4). The location of the proteolysis and the number of cleavages within the APP transmembrane domain by γ-secretase determines the length of final Aβ products and the likelihood of forming plaques.

Of the many transmembrane substrates, processive proteolysis of APP by γ-secretase is the most studied. γ-Secretase first carries out endoproteolytic (ε) cleavage of C99 peptide near the cytosolic end of the transmembrane domain, producing Aβ49 and Aβ48 peptides and their respective AICD co-products (AICD50-99 and AICD49-99, respectively)(5). These initially formed long Aβ peptides are then cut generally every three residues from their C-termini to release tripeptide (and one tetrapeptide) co-products. The two general pathways of γ-secretase processive proteolysis are Aβ48→Aβ45→Aβ42→Aβ38 and Aβ49→Aβ46→Aβ43→Aβ40 (6, 7), producing Aβ42 and Aβ40 as their dominant products, respectively. Among these two, the longer Aβ42 peptide is more prone to aggregate and forms plaques (8). Moreover, early-onset familial AD (FAD) APP mutants can bias the enzyme to produce longer Aβ peptides that are pathological and cause AD (9). The trimming of APP substrate by γ-secretase enzyme is dictated by active site S1’, S2’ and S3’subpockets that respectively bind to P1’, P2’ and P3’ substrate residues (10).

Critical gaps remain in understanding the mechanism of intramembrane processive proteolysis by γ-secretase. Recently reported cryo-EM structures of γ-secretase bound to Notch and APP substrates provided valuable insights into the structural basis of substrate recognition of the enzyme (11, 12). However, artificial structural constraints were included that could affect the enzyme-substrate interactions. Molecular dynamics (MD) simulations have proven useful in understanding the structural dynamics of γ-secretase, notably the enzyme-substrate interactions (13–17). Recently, we computationally restored the wildtype (WT) enzyme-substrate co-structure and applied all-atom simulations using the Gaussian accelerated molecular dynamics (GaMD) method to build the first dynamic model of γ-secretase activation (18). GaMD is an enhanced sampling technique that works by adding a harmonic boost potential to smooth the potential energy surface and reduce system energy barriers (19). Our GaMD simulations captured the extremely slow motions underlying enzyme activation, with the two catalytic aspartates and a coordinated water molecule poised for proteolysis of APP at the ε cleavage site. We showed that the I45F and T48P FAD mutations in APP enhanced the ε cleavage of the amide bond between Leu49-Val50 compared with the WT APP. In contrast, the M51F mutation in APP shifted the ε cleavage to the adjacent Thr48-Leu49 amide bond, changing the proteolysis from the Aβ49 to the Aβ48 pathway. Despite these advances, the detailed atomistic mechanism of processive proteolysis by γ-secretase remains elusive. This is consistent with γ-secretase being a well-known slow-acting enzyme (k_cat_ for APP ε proteolysis ~ 2-6 per hour) (20, 21), making it difficult to capture the dynamic transitions comprising large energy barriers in MD simulations. Hence, despite its importance in the pathogenesis of AD, the mechanism of processive proteolysis (tripeptide trimming) by γ-secretase remains poorly understood.

Here, we report the first dynamic model of tripeptide trimming of Aβ49 to Aβ46 (ζ cleavage) by γ-secretase. Extensive all-atom simulations using a novel Peptide GaMD (Pep-GaMD) method (22) captured the slow dynamic molecular transition from the ε to ζ proteolytic cleavage step. In Pep-GaMD, a boost potential is applied selectively to the essential potential energy of the peptide to effectively model its high flexibility and accelerate its dynamic motions (22). In addition, another boost potential is applied on the protein and solvent to enhance conformational sampling of the protein and facilitate peptide binding. Pep-GaMD has been demonstrated on binding of model peptides to the SH3 protein domains. Independent 1-μs dual-boost Pep-GaMD simulations have captured repetitive peptide dissociation and binding events, which enable calculation of peptide binding thermodynamics and kinetics. The calculated binding free energies and kinetic rate constants agreed very well with the available experimental data (23).

In this study, we have combined biochemical experiments, including matrix-assisted laser desorption/ionization—time-of-flight) mass spectrometry (MALDI-TOF MS), liquid chromatography—tandem mass spectrometry (LC-MS/MS) and western blotting, with Pep-GaMD enhanced sampling simulations to elucidate the mechanism of tripeptide trimming of Aβ49 by γ-secretase. Our findings from Pep-GaMD simulations of WT and five FAD mutants (I45F, A42T, V46F, I45T and T48P) of Aβ49 bound to γ-secretase were highly consistent with quantitative biochemical analysis of their specific proteolytic products, providing important mechanistic insights into tripeptide trimming by the enzyme.

## Results

### Probing ζ cleavage of WT and FAD-mutant Aβ49 by γ-secretase in biochemical experiments

To compare the ζ cleavage of the WT and FAD mutants of APP by γ-secretase, we performed *in vitro* cleavage assay experiments using purified γ-secretase and recombinant APP-based substrate C100-FLAG, which contained the C99 APP C-terminal fragment with an N-terminal start methionine and a C-terminal FLAG epitope tag (24). Efficiency of the cleavage of substrate Aβ49 to products Aβ46 and tripeptide was calculated by measuring Aβ49 production and Aβ49 degradation. To quantify Aβ49 production by ε cleavage of APP substrate, levels of co-products AICD 50-99 were determined using a combination of matrix-assisted laser desorption/ionization time-of-flight mass spectrometry (MALDI-TOF MS) and quantitative western blotting.

First, AICD produced in the assay was immunoprecipitated with anti-FLAG antibodies and detected by MALDI-TOF MS (**Fig. 1A**). For the WT, A42T, V46F and I45T APP substrate, the signal intensities corresponding to AICD 49-99 and AICD 50-99 show higher level of AICD 49-99 than AICD 50-99. However, for mutants I45F and T48P APP substrate, signal intensities show higher level of AICD 50-99 than AICD 49-99. This suggests I45F and T48P favor production of Aβ49 rather than production of Aβ48 while A42T, V46F and I45T favor production of Aβ48 rather than production of Aβ49.

**Figure 1:**
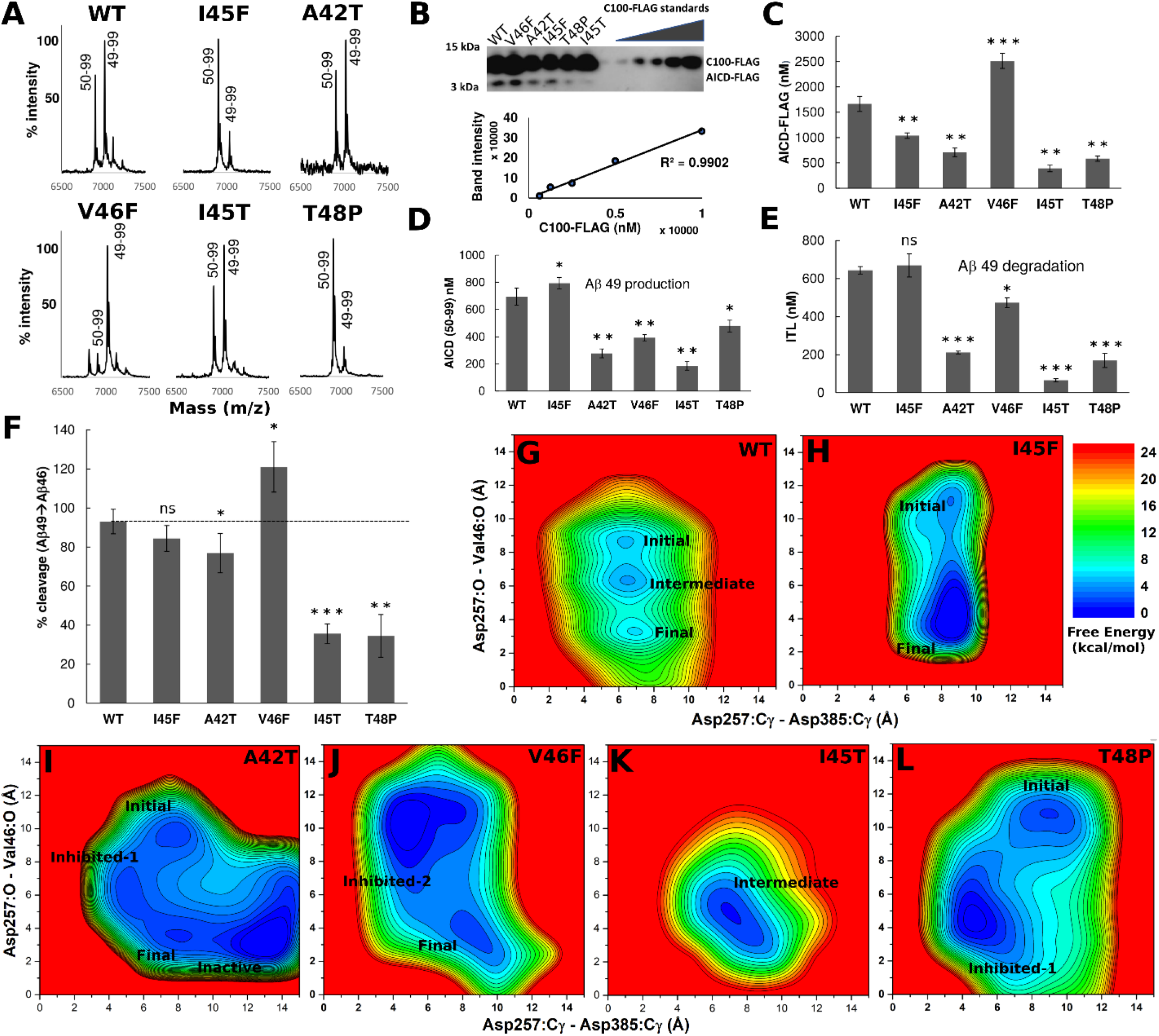
Tripeptide trimming of the wildtype (WT) and FAD mutants of Aβ49 γ-secretase characterized by MS, western blotting and Pep-GaMD simulations. (A) MALDI-TOF MS detection of AICD 50-99 and AICD 49-99 products, (B) Anti-FLAG immunoblot of total AICD-FLAG levels. Purified C100-FLAG at a range of known concentrations was used to generate a standard curve, (C) Quantification of total AICD-FLAG levels from immunoblot by densitometry, (D) Quantification of AICD 50-99 using total AICD levels determined from immunoblot and intensity ratios determined from MALDI-TOF MS, (E) Quantification of ITL tripeptides generated from the first trimming step of WT and FAD mutants of Aβ49, (F) Cleavage efficiency of the first trimming (ζ) step. Grey dotted line denotes cleavage efficiency from WT APP substrate. (G-L) 2D free energy profiles calculated from the Pep-GaMD simulations of (G) WT, (H) I45F, (I) A42T, (J) V46F, (K) I45T and (L) T48P Aβ49 bound to γ-secretase. The distances between the Cγ atoms of Asp257 and Asp385 in PS1 and between the hydroxyl oxygen of PS1 Asp257 and the carbonyl oxygen of Aβ49 Leu49 were selected as the reaction coordinates.

The same reaction mixtures were subjected to quantitative western blotting with anti-FLAG antibodies (**Fig. 1B**), where standards of known concentrations of C100-FLAG were also run to make a standard curve, plotting band intensity against concentration of FLAG-tagged C100. From this standard curve, the concentration of total AICD-FLAG product obtained in the enzyme reaction was quantified (**Fig. 1C**). Quantification of the total AICD revealed increased total AICD production for V46F mutant substrate and decreased total AICD production for A42T, I45F, I45T, and T48P mutant substrates compared to AICD production for the WT. The concentration of AICD 50-99 was calculated using the total AICD level determined by quantitative western blot and the ratio of AICD 49-99 to AICD 50-99 determined from MALDI-TOF MS. (**Fig. 1D**). The calculated concentrations of AICD 50-99 thus provided the level of production of co-product Aβ49. Aβ49 production was slightly increased for I45F mutant, while for all other mutants A42T, V46F, I45T and T48P decreased Aβ49 production was observed compared to Aβ49 production of the WT.

To determine the degradation of Aβ49, we calculated and quantified trimming product tripeptide ITL. The mixtures from the cleavage assay were subjected to LC-MS/MS analysis to detect tripeptides. All substrate constructs studied produced ITL except for T48P mutant which produced IPL due to the replacement of T with P. For quantification of these tripeptides production, standard curves of each peptide were generated by plotting the concentration of synthetic peptide against the integrated areas of the three most abundant ion fragments from MS/MS (**Fig. S1**). The ITL and IPL peptide generated in the γ-secretase cleavage was monitored and quantified. (**Fig. 1E**). The quantification of the trimming product (ITL or IPL) or the Aβ49 degradation reveal decrease in Aβ49 degradation for A42T, V46F, I45T and T48P. For I45F, Aβ49 degradation is similar to that of the WT. Concentration of both Aβ49 production as well as Aβ49 degradation was used to calculate the percent efficiency (**Fig. 1F**). For all constructs, cleavage efficiency was close to 100% except that for two mutants I45T and T48P, the cleavage efficiencies decreased substantially to 35% and 34%, respectively.

### Activation of γ-secretase for tripeptide trimming of Aβ49 was captured in Pep-GaMD simulations

In parallel with the biochemical experiments, Pep-GaMD simulations were carried out on the γ-secretase bound by the WT and the I45F, A42T, V46F, I45T and T48P mutants of Aβ49. The active WT APP-bound γ-secretase was obtained from our previous study (18), and the amide bond between Aβ49 and AICD50-99 was cleaved as the new simulation starting structure **(Fig. S2,** see details in **Methods)**. We initially performed dual-boost GaMD simulations on the γ-secretase bound to Aβ49 with AICD50-99 removed. However, even after running ~6 μs GaMD simulations, we could not effectively sample conformational transitions of the system for ζ cleavage of Aβ49 to Aβ46 **(Fig. S3)**. The distance between the enzyme Asp257 catalytic residue and substrate Val46-Ile47 amide bond presented a computational challenge for conformational sampling, with apparently high energy barriers to overcome. To address the challenge, we applied our recently developed Pep-GaMD (22) method, which selectively boosts the essential potential energy of the peptide to effectively model the peptide flexibility and further improve sampling. We built three different Pep-GaMD simulation systems with γ-secretase bound to Aβ49 in the presence and absence of charged and neutral N-terminal AICD50-99 **(Fig. S4)**. Spontaneous activation of γ-secretase for ζ cleavage of Aβ49 was observed during 600 ns Pep-GaMD simulations with charged N-terminal AICD50-99 **(Figs. S5** and **S6** and **Movie S1)**. The enzyme activation for ζ cleavage was characterized by coordinated hydrogen bonding between the enzyme Asp257 and carbonyl oxygen of substrate Val46. The catalytic aspartates were at a distance of ~7-8 Å between their Cγ atoms, which could accommodate a water molecule for nucleophilic attack of the carbonyl carbon of the scissile amide bond **(Fig S5)**. The water molecule formed hydrogen bonds with both catalytic aspartates and was at ~4 Å distance away from the carbonyl carbon of substrate Val46 residue. The activated γ-secretase conformation was well poised for cleavage of amide bond between Val46 and Ile47 for ζ cleavage of the Aβ49. In the charged N-terminal AICD system, we observed AICD50-99 dissociation in addition to enzyme activation for ζ cleavage **(Fig. S7 and Movie S2)**. The AICD50-99, initially located near the Aβ49, slowly moved downwards to the intracellular PS1 pocket and then dissociated completely from the enzyme. Meanwhile, the AICD50-99 transitioned from β-sheet to a loop/un-structured conformation during the Pep-GaMD simulations **(Fig. S7)**. In comparison, the γ-secretase systems bound to APP in the absence and presence of neutral N-terminal AICD50-99 could not sample enzyme activation for ζ cleavage and AICD50-99 dissociation (**Fig. S6A** and **S6B**). This showed that the presence of charged N-terminal AICD50-99 was crucial for the enzyme activation for ζ cleavage of Aβ49.

Free energy profiles were calculated from Pep-GaMD simulations to characterize the activation of γ-secretase for ζ cleavage of the Aβ49 substrate, for which the distance between the enzyme catalytic aspartates and the distance between the enzyme protonated Asp257 and substrate residue Val46 were selected as reaction coordinates **(Fig. 1G–1L, S6 and S8)**. In the WT Aβ49, three low-energy conformational states were identified from the free energy profile, including “Final”, “Intermediate” and “Initial” **(Figs. 1G, S6C and S9)**. In the “Final” conformational state, the aspartates were ~7-8 Å apart to accommodate the water molecule in between. The substrate Val46 maintained a distance of ~3 Å from the active site Asp257 to form hydrogen bond in the Final active state. In the “Initial” state, the substrate Val46 was distant (~8-9 Å) from the active site Asp257, while the inter-aspartate distance was ~6-7 Å. The “Initial” state represented the active state for the ε cleavage of APP. In the “Intermediate” state, the aspartates remained ~6-7 Å apart, while the Aβ49 peptide (carbonyl oxygen of Val46) was at a distance of ~6 Å from the protonated Asp257 **(Fig. 1G)**.

In the I45F mutant system, two low-energy conformational states, “Initial” and “Final”, were identified from the free energy profile of Pep-GaMD simulations **(Figs. 1H, S6D and S8D and Movie S3)**. Two out of three Pep-GaMD simulations could capture the activation process, as the Asp257 could form stable hydrogen bond with Val46 as reflected in the distance time course plot (**Fig. S6D**). The “Final” state in the free energy profile represented the active conformation of the enzyme for ζ cleavage of the scissile amide bond between Val46 and Ile47 APP residues. In the “Final” conformational state, the two catalytic aspartates were ~7-8 Å apart, and APP Val46 was ~3 Å distance away from the protonated aspartate. In the “Initial” state, the substrate Val46 was further away from the catalytic aspartate (~6-7 Å), and the aspartates were ~7-8 Å distance away from each other **(Fig. 1H)**.

In the A42T mutant APP system, four low-energy conformational states were identified from the free energy profile **(Figs. 1I, S6E and S8E and Movie S4)**. Mutation of Ala42 to Thr42 caused the enzyme-substrate complex to be more flexible and sample a larger conformational space. In addition to the “Initial” and “Final” states, two new “Inhibited-1” and “Inactive” conformational states were identified for the A42T mutant system. The catalytic aspartates were ~4-5 Å (too close) apart in the “Inhibited-1” state and 13 Å away (too far) in the “Inactive” state. In the “Inhibited” state, the catalytic aspartates could not accommodate a water molecule between them and hence was inhibited from proteolytic activation. APP Val46 was ~4-5 Å from the protonated Asp257 in this “Inhibited-1” state. In the “Inactive” state, the aspartates were ~13 Å apart and thus too far to form the dual hydrogen bonds with the water in between them, even though the Asp257 could form a hydrogen bond with the Val46 carbonyl oxygen. This hindered activation required for ζ cleavage.

In the V46F mutant system, two low-energy conformational states were identified, including “Inhibited-2” and “Final” **(Figs. 1J, S6F and S8F and Movie S5)**. Like other γ-secretase systems, the “Final” state corresponded to the active conformation of the enzyme poised for ζ cleavage of Aβ49. Moreover, the “Inhibited-2” state had the two aspartates at proximity (~4-5 Å) between the Cγ atoms and unable to accommodate a water molecule in between for enzyme activation. APP substrate was ~10 Å away from active site Asp257 in the “Inhibited-2” state.

Furthermore, Pep-GaMD simulations were carried out on I45T and T48P mutant Aβ49-bound γ-secretase **(Figs. 1K–1L, S6G-S6H and S8G-S8H)**. Both of these mutant systems were not able to activate the enzyme for ζ cleavage, being consistent with the experimental results where the ζ cleavage efficiency dropped to about one third compared to that of the WT. In the Pep-GaMD free energy profile of the I45T mutant system, only one “Intermediate” low-energy conformational state was identified. This “Intermediate” state was the same low-energy conformation as the one in the WT system. For the T48P system, the hydrogen bond between APP Val46 and the protonated Asp257 was formed for a certain time in one of the three Pep-GaMD production simulations **(Fig. S6H)**. However, in the free energy profile, we could identify two low-energy conformational states, including “Initial” and “Inhibited-1”, but not the “Final” active state **(Fig. 1L)**. The “Inhibited-1” state resembled the one identified in the A42T mutant system. The “Initial” conformational state was the same as the one identified in the WT, I45F and A42T systems. These Pep-GaMD simulation findings were consistent with the biochemical experiments, verifying the I45T and T48P systems as negative controls.

### Conformational changes in activation of γ-secretase for tripeptide trimming of Aβ49

We calculated root mean square fluctuations (RMSFs) of γ-secretase bound by the WT and FAD-mutant APP from Pep-GaMD simulations **(Fig. S10 and Movie S6)**. In the WT Aβ49-bound γ-secretase, the TM2, TM6, TM6a and C-terminus of TM9 helix were flexible in the catalytic PS1 subunit. The Pen-2 subunit exhibited high fluctuations with ~ 3 Å RMSF. Helices a1, a2, a5, a12, a17, and TM domain of nicastrin were also flexible during the Pep-GaMD simulations. Structural clustering was performed on Pep-GaMD snapshots of the system using hierarchical agglomerative algorithm in CPPTRAJ (25) (see **Methods**). The top-ranked cluster was selected as the representative “Final” active conformation for the ζ cleavage of Aβ49. The starting structure from ε cleavage of APP was obtained as the “Initial” active conformation. The catalytic PS1 of the “Final” conformation was compared to that of the “Initial” conformation in **Fig. 2A**. Relative to the “Initial” conformation, the substrate helical domain tilted by ~50 degrees in the “Final” conformation **(Figs. 2A** and **2B)**. Residue Leu49 in the substrate C-terminus moved downwards by ~5 Å **(Figs. 2B**, **4A** and **S11)**. The last residue in a helical conformation in the “Final” state of Aβ49 was Thr43 whereas it was Ile45 in the “Initial” state. In transition from the “Initial to the “Final” conformational state, two substrate residues, Val44 and Ile45 unwound their helical conformation and changed to a turn/loop conformation. Residues Thr43 and Ile45 were in similar positions in the “Initial” and “Final” active conformations relative to the membrane perpendicular axis (**Fig. S11**). In comparison, the substrate C-terminal Leu49 moved downwards by ~5 Å while straightening the C-terminal loop **(Figs. 2B and S11B)**.

**Figure 2:**
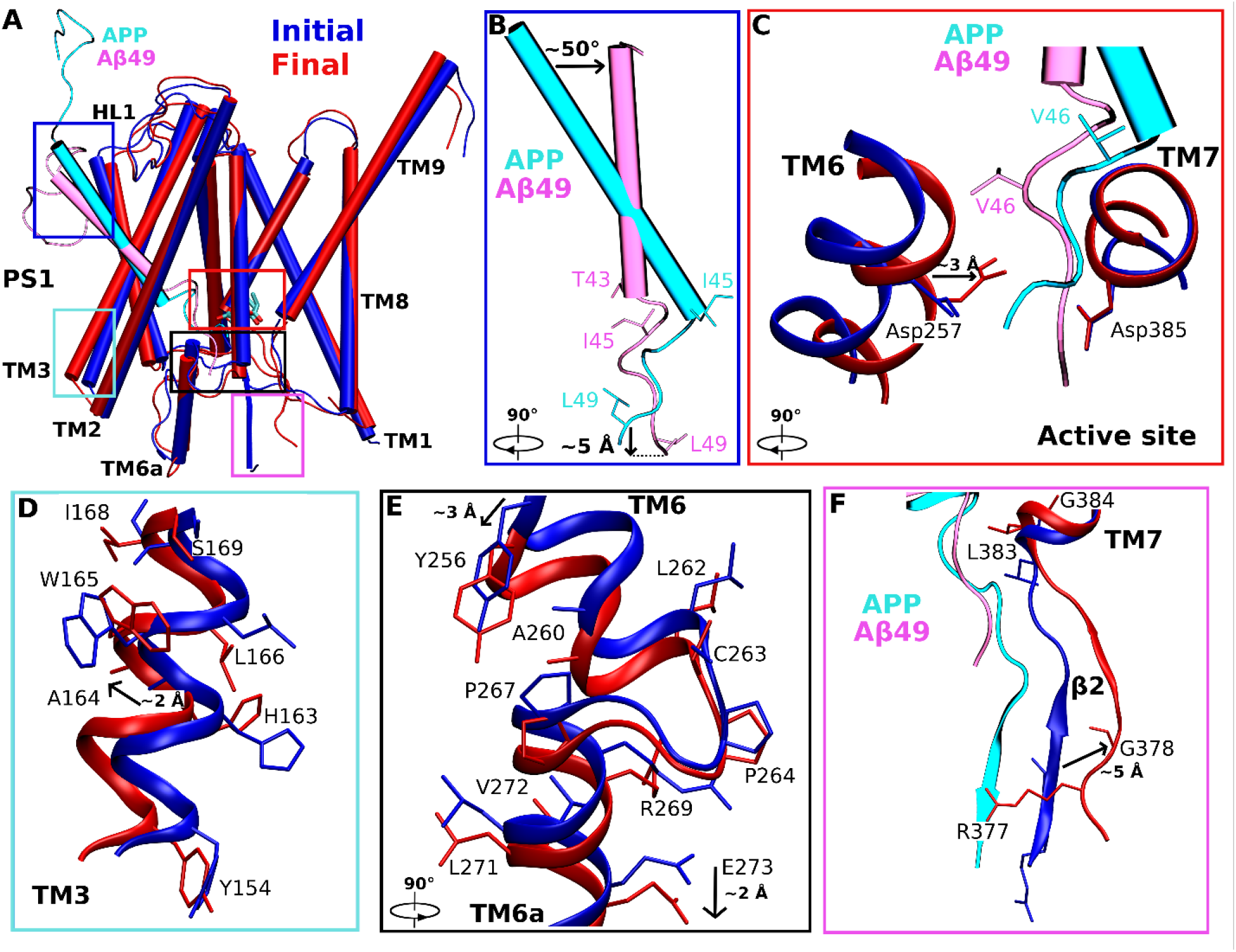
Conformational changes of the PS1 catalytic subunit and substrate during activation of γ-secretase for tripeptide trimming of Aβ49 in Pep-GaMD simulations. (A) Comparison of the Initial (active for ε cleavage, blue) and Final (active for ζ cleavage, red) conformations of the Aβ49-bound PS1. The enzyme activation for tripeptide trimming was characterized by coordinated hydrogen bonding between the enzyme Asp257, carbonyl oxygen of Aβ49 Val46 and a water molecule accommodated between the two aspartates poised for cleavage of the amide bond between Val46 and Ile47 residues. (B-F) Conformational changes of (B) Aβ49 substrate, (C) catalytic aspartates, (D) TM3, (E) TM6 and TM6a, and (F) β2 strand from the Initial to the Final conformational state. The helical domain of Aβ49 tilted by ~50° and residue Leu49 at the C-terminus of Aβ49 moved downwards by ~5 Å. Protonated catalytic Asp257 moved ~3 Å towards the Aβ49 substrate. The enzyme TM3 moved outwards by ~2 Å and TM6a moved downwards by ~2 Å. The enzyme β2 strand (N-terminus of TM7) moved away from APP and closer towards the β1 strand (C-terminus of TM6a) by ~5 Å.

**Figure 3:**
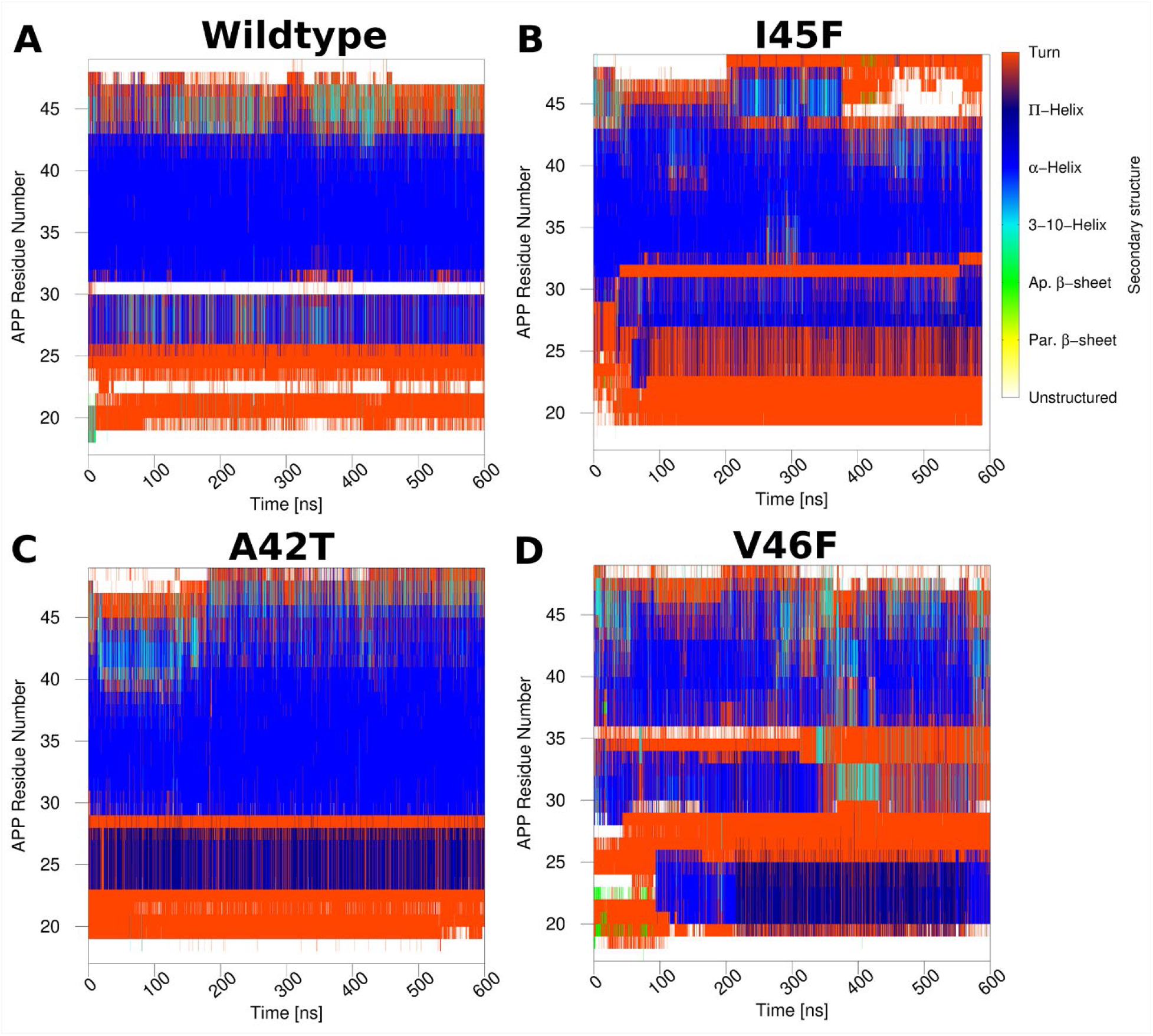
Time-dependent secondary structures of Aβ49 bound to γ-secretase calculated from the Pep-GaMD simulations. (A) WT, (B) I45F, (C) A42T and (D) V46F systems of Aβ49. Results from other simulations are plotted in **Figs. S12 and S15**.

**Figure 4:**
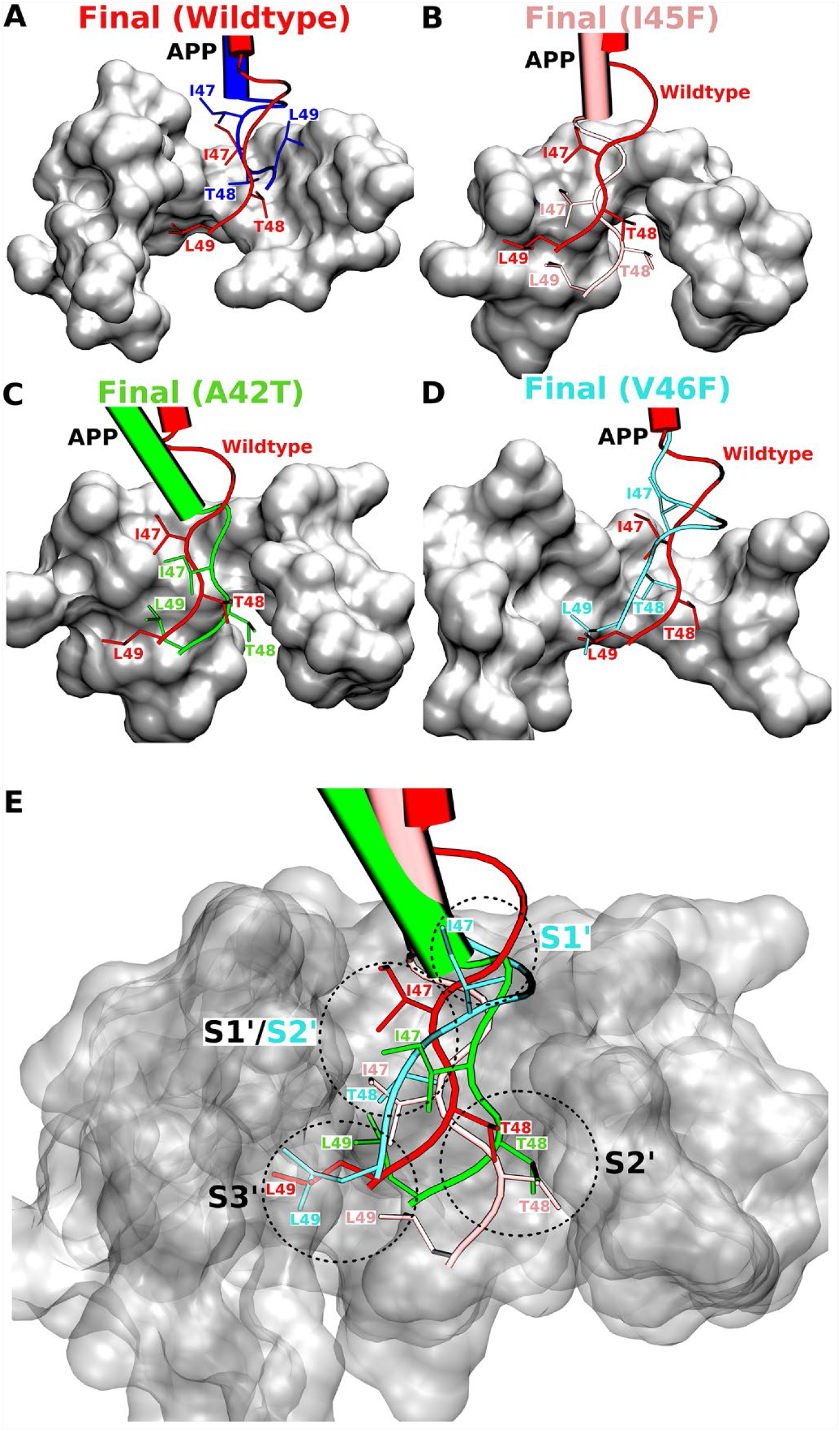
Active-site conformations of γ-secretase for tripeptide trimming of Aβ49 observed in the Pep-GaMD simulations. (A-D) Conformations of the substrate P’, P2’ and P3’ residues in the Final active conformations of the (A) WT (red), (B) I45F (pink), (C) A42T (green) and (D) V46F (cyan) Aβ49 systems. (E) Comparison of the PS1 active-site S1’, S2’ and S3’ pockets that accommodate the WT and mutants of Aβ49.

At the enzyme active site, the catalytic Asp385 did not have significant movement during the adjustments for substrate peptide trimming (**Fig. 2C**). In comparison, the protonated catalytic Asp257 moved by ~3 Å towards the substrate. Asp257 moved forward to form a hydrogen bond with the carbonyl oxygen of the scissile amide bond between the substrate residues Val46 and Ile47. Similarly, TM3 moved outwards by ~2 Å **(Fig. 2D)**, and TM6a moved downwards by ~2 Å **(Fig. 2E)**. Flexibility in these helices involved important FAD mutation sites including Tyr154, His163, Ala164, Leu166, Trp165, Ser169, Ile168, Tyr256, Ala260, Leu262, Cys263, Pro264, Pro267, Arg269, Val272 and Leu271 (www.alzforum.org). Trp165 and His163 from TM3 and Arg269 from TM6a showed significant movements in their side chains. With a major part of C-terminus of APP absent (as AICD dissociates, see next section) the β2 loop at N-terminus of TM7 moved away from the APP by ~5 Å in the Final state as compared to the Initial state **(Fig. 2F)**. FAD mutation residues in the β2-TM7 region including Arg377, G378, L383 and G384 showed flexibility in the simulations. In particular, residue Arg377 reoriented its side chain in the “Final” conformational state.

### Changes in secondary structures of the WT and FAD-mutant Aβ49 during tripeptide trimming

Secondary structures of the WT and FAD-mutant Aβ49 bound to γ-secretase were recorded during the Pep-GaMD simulations and plotted in **Figs. 3 and S12**. Changes in secondary structures of Aβ49 during ζ cleavage were compared to that of APP substrate (“Initial” active conformation) during ε cleavage from our previous study (18) **(Fig. S13)**. Unwinding of the helix C-terminus in Aβ49 during ζ cleavage was observed in the secondary structure plot. During the ε cleavage, the C-terminus of the WT APP substrate could maintain helical conformation up to Ile45/Val46 **(Fig. S13)**. In comparison, WT Aβ49 was helical up to Thr43 in the C-terminal region **(Figs. 3A and 2B)**. About 2-3 residues unwound near the ζ cleavage site to expose the scissile amide bond between Val46 and Ile47 to the catalytic aspartates and the coordinated water for activation. A new helix was formed for residues Ser26 to Ala30 in the Aβ49 during the transition from ε to ζ cleavage in the WT system **(Figs. 3A and S14)**. With the 50° tilt of Aβ49 peptide in the space between TM2 and TM3, the N-terminus is exposed to the hydrophobic lipid bilayer **(Fig. S14)**. This helped the N-terminal loop to transition to a α-helical conformation. The effects of the mutations on the new helical conformation is mentioned and explained in the next paragraph. A turn/unstructured conformation at residues Ala30-Ile31 separated these two helices. In addition, the N-terminus of Aβ49 lost its interactions with the hydrophobic loop 1 (HL1) because of the tilting away from this loop.

Similarly, secondary structural changes were recorded for the I45F, A42T and V46F Aβ49 mutant systems **(Figs. 3B–3D and S15-S16)**. Like the WT, the I45F and A42T Aβ49 mutants maintained a helical conformation up to Thr43 at the N-terminus during the Pep-GaMD simulations. C-terminal residues after the Thr43, which included the ζ cleavage site bond between Val46 and Ile47, were observed mostly in a turn/unstructured conformation. This allowed the catalytic aspartates and water to approach the scissile amide bond for forming coordinated hydrogen bonds required for this cleavage. Likewise, bands of new helix formation were observed in the secondary structure plots from Asn27 to Ile31 and from Asp23 to Lys28 for I45F- and A42T-mutant Aβ49 systems, respectively **(Fig. 3B and 3C)**. The new helix formed was due to its exposure to the hydrophobic lipid membrane. V46F Aβ49 was observed to be the most dynamic in terms of secondary structure changes **(Fig. 3D)**. A band of helix was observed between Gly29 to Thr43, with a turn conformation formed between at Leu34 – Val36. Thr43 to Ile47 transitioned between helix and turn conformations during the Pep-GaMD simulations of the V46F Aβ49. Like the WT and other mutant systems, new N-terminal helix formation was observed at residues Phe20 to Gly25 in the V46F mutant APP. The hydrophobic lipid environment helped these residues transition from turn to a helical conformation in the V46F mutant APP.

### Active-site subpockets formed in γ-secretase for tripeptide trimming

The “Final” active conformational state of Aβ49-bound γ-secretase was further analyzed for the P1’, P2’ and P3’ substrate residues at the ζ cleavage active site and the respective S1’, S2’ and S3’ subpockets in which they reside (10). The S1’ subpocket accommodating the P1’ residue in the WT Aβ49 was formed by residues from PS1 TM6a helix, β1 loop, TM3 helix and TM7 N-terminal region **(Figs. 2 and 4A)**. The residues that formed the subpockets are listed in **Table S1**. The S2’ subpocket occupied by the P2’ substrate residue consisted of residues from PS1 TM6 helix, TM6a helix, PAL motif of TM9 helix, β1 and β2 loop region. Moreover, the S3’ subpocket accommodating the P3’ residue was formed by residues from PS1 TM6 helix, TM6a helix and β1 loop. In reference to Aβ49, S1’ and S3’ pockets were located on the same side (TM6a and TM3 helices) whereas the S2’ pocket was located on the opposite side (TM6 and TM9 helices).

Similarly, the “Final” active conformational states of the I45F and A42T mutant Aβ49-bound γ-secretase systems had the same subpockets formed at the active site during the ζ cleavage as that of the WT system **(Fig. 4B–4C and Table S1)**. In the I45F and A42T “Final” active conformation, the S1’ and S3’ subpockets occupied by the respective P1’ and P3’ substrate residues consisted of residues from PS1 TM6 helix, TM6a helix, TM7 helix, β1 and β2 loop region. In comparison, the S2’ subpocket was located on the opposite side of Aβ49 and consisted of residues from PS1 TM6 helix, TM6a helix, PAL motif of TM9 helix, β1 and β2 loop. Furthermore, in the V46F “Final” active conformation, the locations of the S1’ and S2’ subpockets accommodating P1’ and P2’ Aβ49 substrate residues, respectively, were different as compared to that of the WT system **(Fig. 4C-D)**. The S1’ pocket occupied by the P1’ residue of Aβ49 consisted of residues from TM6 helix, TM6a helix and TM2 helix **(Table S1)**. The S2’ subpocket occupied by the P2’ residue of the V46F mutant in the “Final” active state was the same as the S1’ subpocket in the “Final” active state of the WT, I45F and A42T systems **(Fig. 4C-D)**. Moreover, the S3’ subpocket accommodating the P3’ substrate residue in the V46F mutant was the same as the one of the WT, I45F and A42T systems (**Fig. 4E**).

## Discussion

Current AD treatments ease symptoms, but none has been clearly demonstrated to slow or halt disease progression. While the molecular cause of AD remains poorly understood, the hallmark pathological criteria for AD diagnosis is the deposition of amyloid-β (Aβ) plaques in the brain (26). Aβ peptides are products of processive proteolysis by γ-secretase. Dominant missense mutations in the substrate (APP) and the enzyme (presenilin component of γ-secretase) cause early-onset FAD, and these mutations result in deficient carboxypeptidase trimming of initially formed long Aβ peptides to shorter secreted forms (27–29). Yet the mechanism of processive proteolysis of APP by γ-secretase is unknown. Recent reports of cryo-EM structures of γ-secretase bound to APP and Notch substrates as well as to γ-secretase inhibitors and modulators revealed details of the structural basis of substrate recognition as well as enzyme inhibition and modulation (11, 12, 30). Regardless, static conformations of the enzyme cannot explain the underlying mechanism of enzyme activation and substrate processing. Essentially nothing is known about the dynamic mechanism of processive proteolysis by γ-secretase.

Here, we have addressed the issue by combining novel Pep-GaMD enhanced sampling simulations and biochemical experiments. Different systems of γ-secretase bound by the WT and FAD-mutant Aβ49 substrates were investigated to understand the first tripeptide trimming step, ζ cleavage **(Fig. 5)**. Four γ-secretase systems—bound to the WT, I45F, A42T and V46F Aβ49 in presence of charged N-terminal AICD—underwent activation for ζ cleavage during 600 ns Pep-GaMD simulations **(Fig. 5B)**. This was consistent with biochemical experiments, as these mutant systems showed similar efficiencies for the Aβ49 to Aβ46 proteolytic step (ζ cleavage). In comparison, γ-secretase bound by I45T and T48P Aβ49 showed little or no sample activation **(Fig. 5C)**. Furthermore, Aβ49-bound γ-secretase in the absence of AICD was not able to sample the “Final” active state for ζ cleavage of the substrate **(Fig. S17A)**, similarly for γ-secretase bound to WT Aβ49 in the presence of neutral N-terminal AICD **(Fig. S17B)**. This highlighted the importance of AICD and its N-terminal charge in facilitating processive proteolysis by γ-secretase. Following ε cleavage, both the C-terminus of Aβ49 and N-terminus of AICD at the active site could be exposed to water molecules and thus charged at physiological pH 7 (as carboxylate and ammonium, respectively). The charged state likely aided movement toward the polar aqueous environment and away from the hydrophobic transmembrane interior of the PS1 active site. Indeed, the AICD with charged N-terminus could dissociate from PS1 in the Pep-GaMD simulations, while the charged C-terminus of Aβ49 helped prepare for the next cleavage during processive proteolysis by γ-secretase.

**Figure 5:**
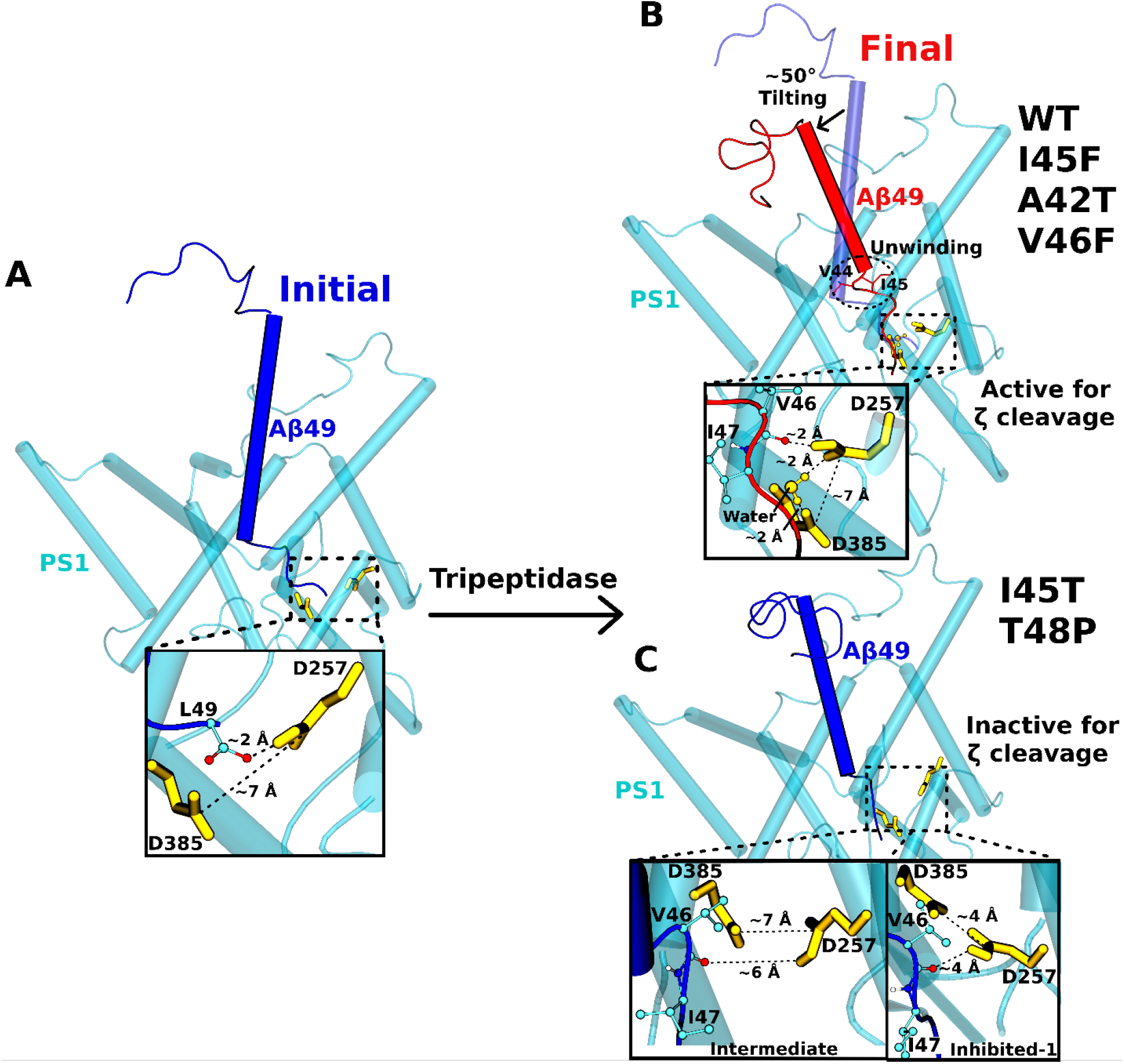
Dynamic model of tripeptide trimming by γ-secretase. (A) The “Initial” conformational state of Aβ49 bound γ-secretase. (B) The WT Aβ49 and its I45F, A42T and V46F mutants were able to transition to the “Final” state with ~50° tilting of the helical domain and unwinding of the helix C-terminus (residues V44-I45) and became poised for ζ cleavage of the V46-I47 amide bond by γ-secretase. (C) In contrast, the I45T and T48P mutant Aβ49-bound γ-secretase were trapped in the “Intermediate” or “Inhibited-1” state, being inactive for ζ cleavage of the substrate.

During ζ cleavage activation, two residues unwound from the C-terminus of the Aβ49 helix, changing to a turn conformation **(Fig. 5B)**. This was observed in the time courses of the substrate secondary structures as well. Unlike the helical conformation, the loop/turn conformation facilitated exposure of the scissile amide bond to the catalytic aspartates and the coordinated water molecule. In parallel, positions of the Thr43 and Ile45 residues in the “Initial” and “Final” states relative to the membrane were similar, whereas the C-terminal residue Leu49 moved downwards by ~5 Å. Moreover, the helical domain of Aβ49 tilted by ~50 degrees **(Fig. 5B)**. Thus, tilting of the helical domain and unwinding of C-terminal helix in the substrate apparently facilitated the proteolytic progression from ε to ζ cleavage by γ-secretase. Helix unwinding was accompanied by straightening of the C-terminal loop/turn and downward movement of the terminal residue Leu49. Similarly, the β-sheet conformation between the APP C-terminus and the β1 loop was broken as ε cleavage product AICD50-99 dissociates. This caused the β1 loop to move away from the APP C-terminus by ~5 Å. This region has been suggested to be important for substrate recognition and proteolytic processing (12). Similarly, γ-secretase inhibitors (GSIs) and transition state analogs (TSAs) bind to this region (30). The present study also shows an important role of this region in activation of γ-secretase for ζ cleavage of Aβ49.

Pep-GaMD captured the enzyme activation for ζ cleavage for γ-secretase systems bound to WT and three FAD-mutant (I45F, A42T and V46F) APP substrates. The low-energy “Final” active conformation was identified in the Pep-GaMD free energy profiles of all these systems. However, the PMF profiles representing each enzyme systems was different in terms of distinct low-energy states and the conformational space sampled by the enzyme during ζ cleavage. The I45F and V46F mutant systems sampled two low-energy conformations with the I45F system being the least conformationally dynamic **(Fig. 1H, J)**. Three and four low-energy states were identified from free energy profiles of the WT and the A42T mutant systems, respectively, with the A42T mutant system being more dynamic **(Fig. 1G, I)**. Each system had its own set of conformations and a distinct activation pathway. This suggested that the enzyme is remarkably dynamic, consistent with its ability to cleave over 100 different substrates (31).

In the “Final” active state of γ-secretase poised for the ζ cleavage, subpockets were formed in the active site that were different from that formed for the ε cleavage **(Fig. 4 and S18)**. This finding was consistent with the observation that the C-terminus of Aβ49 during ζ cleavage did not form a β-sheet conformation with the PS1 TM6a β2 region, instead adopting a loop conformation **(Fig. 2F)**. The locations of the active site subpockets formed for ζ cleavage were compared to those formed for ε cleavage **(Fig. S18)**. Moreover, the locations of S1’, S2’ and S3’ subpockets formed for ζ cleavage were the same for different γ-secretase systems bound to the WT and mutant Aβ49 except for the V46F mutant system. The S2’ subpocket accommodating the T48 P2’ residue formed for ζ cleavage was the same as the S1’ subpocket accommodating the V50 P1’ subpocket formed for ε cleavage. In contrast, the ζ cleavage S3’ subpocket for the L49 P3’ residue was the same as the ε cleavage S3’ subpocket for the L52 P3’ residue. The S1’ subpocket for the I47 P1’ residue for ζ cleavage and the S2’ subpocket for the M51 P2’ residue for ε cleavage had their own unique location in their respective Final active states. Regardless, S1’/S3’ and S2’ subpockets were located on opposite sides of the substrate in both of the “Initial” and “Final” active states.

In summary, the Pep-GaMD simulation findings were highly consistent with mutagenesis and mass spectrometry experiments, which quantified AICD50-99 (and therefore Aβ49) production as well as the ζ-cleaved tripeptide product of these APP FAD-mutant systems. Overall, our combined Pep-GaMD simulations and biochemical experiments have revealed the dynamic mechanism of tripeptide trimming by γ-secretase.

## Materials and Method Summary

γ-Secretase assays were performed with purified γ-secretase and C100-FLAG substrates and analyzed previously reported (20). Briefly, the reaction mixture was subjected to MALDI-TOF mass spectrometry to detect AICD-FLAG species. The same reaction mixture was western blotted with anti-Flag antibodies along with C100-FLAG standard to quantify the total AICD species. The reaction was also subjected to LC-MS/MS to detect tripeptides generated in the reactions. Pep-GaMD simulations was performed on the γ-secretase activation for ζ cleavage of Aβ49. Active APP-bound γ-secretase was taken from the previous study (18) and the amide bond between Aβ49 and AICD50-99 was cleaved as the starting structure. The simulation systems of γ-secretase bound by the WT and the I45F, A42T, V46F, I45T and T48P mutants of APP are summarized in **Table 1**. The CHARMM36m (32) parameter set was used for the protein and POPC lipids. Initial energy minimization and thermalization of the γ-secretase complex followed the same protocol as used in the previous GaMD simulations of membrane proteins (33, 34). The Pep-GaMD implemented in the GPU version AMBER 20 (32) was then applied to perform the simulations using the dual-boost scheme, which included 10 ns short cMD simulation used to collect potential statistics for calculating the Pep-GaMD acceleration parameters, 50 ns equilibration after adding the boost potential and finally multiple independent 600 ns production runs with randomized initial atomic velocities. The simulation frames were saved every 0.2 ps for analysis. The VMD (35) and CPPTRAJ (25) tools were used to visualize and analyze the Pep-GaMD trajectories. The *PyReweighting* toolkit (36) was applied to reweight Pep-GaMD simulations for free energy calculations by combining all simulation trajectories for each system. Details of the γ-secretase assays, mass spectrometry, Western blotting, Pep-GaMD method, system setup, simulation protocol, energetic reweighting, and simulation analysis are provided in **Supporting Information**.

**Table 1:**
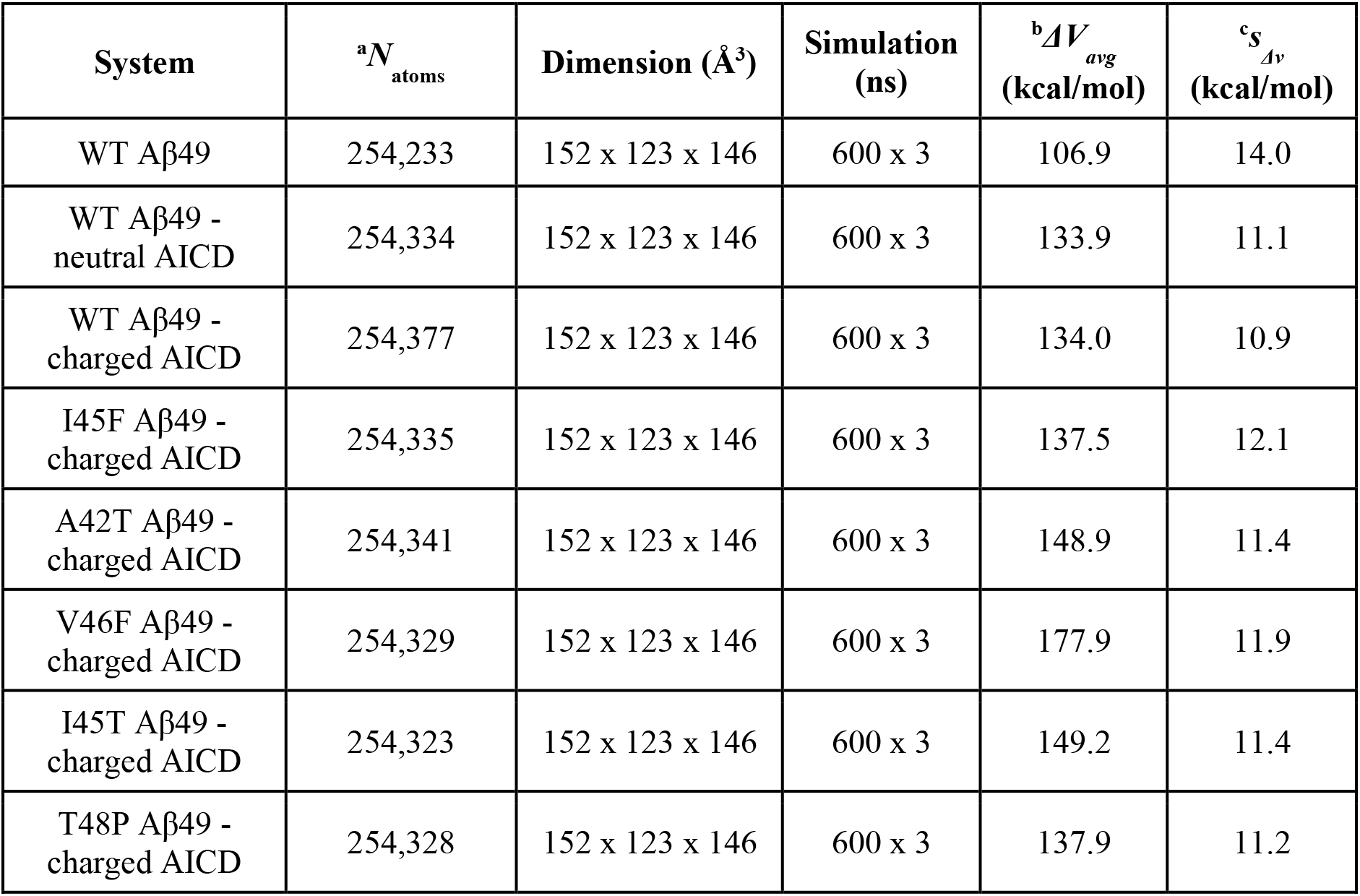
Summary of Pep-GaMD simulations performed on different systems of γ-secretase bound by Aβ49 and AICD50-99 peptides. ^a^*N*_atoms_ is the total number of atoms in the simulation systems. ^b^Δ*V_avg_* and ^c^*σ*_Δ*V*_ are the average and standard deviation of the Pep-GaMD boost potential, respectively.

| System | $^a N_{\text{atoms}}$ | Dimension ($\text{\AA}^3$) | Simulation (ns) | $^b \Delta V_{\text{avg}}$ (kcal/mol) | $^c \sigma_{\Delta V}$ (kcal/mol) |
| --- | --- | --- | --- | --- | --- |
| WT Aβ49 | 254,233 | 152 x 123 x 146 | 600 x 3 | 106.9 | 14.0 |
| WT Aβ49 - neutral AICD | 254,334 | 152 x 123 x 146 | 600 x 3 | 133.9 | 11.1 |
| WT Aβ49 - charged AICD | 254,377 | 152 x 123 x 146 | 600 x 3 | 134.0 | 10.9 |
| I45F Aβ49 - charged AICD | 254,335 | 152 x 123 x 146 | 600 x 3 | 137.5 | 12.1 |
| A42T Aβ49 - charged AICD | 254,341 | 152 x 123 x 146 | 600 x 3 | 148.9 | 11.4 |
| V46F Aβ49 - charged AICD | 254,329 | 152 x 123 x 146 | 600 x 3 | 177.9 | 11.9 |
| I45T Aβ49 - charged AICD | 254,323 | 152 x 123 x 146 | 600 x 3 | 149.2 | 11.4 |
| T48P Aβ49 - charged AICD | 254,328 | 152 x 123 x 146 | 600 x 3 | 137.9 | 11.2 |

## Data Availability

All data are included in the article and/or *Supporting Information*.

## Acknowledgements

This work used supercomputing resources with allocation award TG-MCB180049 through the Extreme Science and Engineering Discovery Environment (XSEDE), which is supported by National Science Foundation grant number ACI-1548562, and project M2874 through the National Energy Research Scientific Computing Center (NERSC), which is a U.S. Department of Energy Office of Science User Facility operated under Contract No. DE-AC02-05CH11231, and the Research Computing Cluster at the University of Kansas. This work was supported in part by the startup funding in the College of Liberal Arts and Sciences at the University of Kansas (Y.M.) and GM122894 from the National Institutes of Health (M.S.W.).

## Author Contributions

A.B., M.S.W. and Y.M. designed research; A.B. and S.D. performed research; A.B., S.D., H.N.D., J.W., S.B., M.S.W. and Y.M. analyzed data: A.B., S.D., M.S.W. and Y.M. wrote the manuscript.

## References

1. J. Weller, A. Budson, Current understanding of Alzheimer’s disease diagnosis and treatment. F1000Research 7 (2018).

2. S. Funamoto, S. Tagami, M. Okochi, M. Morishima-Kawashima (2020) Successive cleavage of β-amyloid precursor protein by γ-secretase. in Seminars in cell & developmental biology (Elsevier), pp 64–74.

3. M. L. Hemming, J. E. Elias, S. P. Gygi, D. J. Selkoe, Proteomic Profiling of gamma-Secretase Substrates and Mapping of Substrate Requirements. Plos Biol 6, 2314–2328 (2008).

4. G. Guner, S. F. Lichtenthaler, The substrate repertoire of gamma-secretase/presenilin. Semin Cell Dev Biol 105, 27–42 (2020).

5. Y. Gu et al., Distinct intramembrane cleavage of the β-amyloid precursor protein family resembling γ-secretase-like cleavage of Notch. Journal of Biological Chemistry 276, 35235–35238 (2001).

6. Y. Qi-Takahara et al., Longer forms of amyloid beta protein: implications for the mechanism of intramembrane cleavage by gamma-secretase. J Neurosci 25, 436–445 (2005).

7. M. Takami et al., gamma-Secretase: successive tripeptide and tetrapeptide release from the transmembrane domain of beta-carboxyl terminal fragment. J Neurosci 29, 13042–13052 (2009).

8. D. J. Selkoe, J. Hardy, The amyloid hypothesis of Alzheimer’s disease at 25 years. EMBO molecular medicine 8, 595–608 (2016).

9. R. E. Tanzi, The genetics of Alzheimer disease. Cold Spring Harbor perspectives in medicine 2, a006296 (2012).

10. D. M. Bolduc, D. R. Montagna, M. C. Seghers, M. S. Wolfe, D. J. Selkoe, The amyloid-beta forming tripeptide cleavage mechanism of γ-secretase. Elife 5, e17578 (2016).

11. G. Yang et al., Structural basis of Notch recognition by human γ-secretase. Nature 565, 192–197 (2019).

12. R. Zhou et al., Recognition of the amyloid precursor protein by human γ-secretase. Science 363 (2019).

13. M. Hitzenberger, M. Zacharias, Structural Modeling of gamma-Secretase A beta(n) Complex Formation and Substrate Processing. ACS chemical neuroscience 10, 1826–1840 (2019).

14. A. Götz et al., Modulating hinge flexibility in the APP transmembrane domain alters γ-secretase cleavage. Biophysical journal 116, 2103–2120 (2019).

15. B. Dehury, N. Tang, K. P. Kepp, Molecular dynamics of C99-bound γ-secretase reveal two binding modes with distinct compactness, stability, and active-site retention: implications for Aβ production. Biochem J 476, 1173–1189 (2019).

16. A. K. Somavarapu, K. P. Kepp, Membrane dynamics of γ-secretase provides a molecular basis for β-amyloid binding and processing. ACS chemical neuroscience 8, 2424–2436 (2017).

17. A. Götz et al., Increased H-Bond Stability Relates to Altered ε-Cleavage Efficiency and Aβ Levels in the I45T Familial Alzheimer’s Disease Mutant of APP. Sci Rep-Uk 9, 1–12 (2019).

18. A. Bhattarai, S. Devkota, S. Bhattarai, M. S. Wolfe, Y. Miao, Mechanisms of γ-Secretase Activation and Substrate Processing. ACS Central Science (2020).

19. Y. Miao, V. A. Feher, J. A. McCammon, Gaussian Accelerated Molecular Dynamics: Unconstrained Enhanced Sampling and Free Energy Calculation. J Chem Theory Comput 11, 3584–3595 (2015).

20. S. Devkota, T. D. Williams, M. S. Wolfe, Familial Alzheimer’s disease mutations in amyloid protein precursor alter proteolysis by γ-secretase to increase amyloid β-peptides of≥ 45 residues. Journal of Biological Chemistry 296, 100281 (2021).

21. F. Kamp et al., Intramembrane proteolysis of β-amyloid precursor protein by γ-secretase is an unusually slow process. Biophysical journal 108, 1229–1237 (2015).

22. J. Wang, Y. Miao, Peptide Gaussian accelerated molecular dynamics (Pep-GaMD): Enhanced sampling and free energy and kinetics calculations of peptide binding. The Journal of Chemical Physics 153, 154109 (2020).

23. Y. Xue, T. Yuwen, F. Zhu, N. R. Skrynnikov, Role of electrostatic interactions in binding of peptides and intrinsically disordered proteins to their folded targets. 1. NMR and MD characterization of the complex between the c-Crk N-SH3 domain and the peptide Sos. Biochemistry 53, 6473–6495 (2014).

24. Y.-M. Li et al., Presenilin 1 is linked with γ-secretase activity in the detergent solubilized state. Proceedings of the National Academy of Sciences 97, 6138–6143 (2000).

25. D. R. Roe, T. E. Cheatham, PTRAJ and CPPTRAJ: Software for Processing and Analysis of Molecular Dynamics Trajectory Data. J Chem Theory Comput 9, 3084–3095 (2013).

26. B. Dubois et al., Advancing research diagnostic criteria for Alzheimer’s disease: the IWG-2 criteria. The Lancet Neurology 13, 614–629 (2014).

27. O. Quintero-Monzon et al., Dissociation between the Processivity and Total Activity of gamma-Secretase: Implications for the Mechanism of Alzheimer’s Disease-Causing Presenilin Mutations. Biochemistry-Us 50, 9023–9035 (2011).

28. M. A. Fernandez, J. A. Klutkowski, T. Freret, M. S. Wolfe, Alzheimer Presenilin-1 Mutations Dramatically Reduce Trimming of Long Amyloid beta-Peptides (A beta) by gamma-Secretase to Increase 42-to-40-Residue A beta. Journal of Biological Chemistry 289, 31043–31052 (2014).

29. S. Devkota, T. D. Williams, M. S. Wolfe, Familial Alzheimer’s disease mutations in amyloid protein precursor alter proteolysis by gamma-secretase to increase amyloid beta-peptides of >/=45 residues. J Biol Chem 296, 100281 (2021).

30. G. Yang et al., Structural basis of gamma-secretase inhibition and modulation by small molecule drugs. Cell 184, 521–533 e514 (2021).

31. G. Güner, S. F. Lichtenthaler (2020) The substrate repertoire of γ-secretase/presenilin. in Seminars in Cell & Developmental Biology (Elsevier).

32. D. A. Case et al., Amber 2020. (2020).

33. K. Kappel, Y. Miao, J. A. McCammon, Accelerated Molecular Dynamics Simulations of Ligand Binding to a Muscarinic G-protein Coupled Receptor. Quarterly Reviews of Biophysics 48, 479–487 (2015).

34. Y. Miao, J. A. McCammon, Gaussian Accelerated Molecular Dynamics: Theory, Implementation and Applications. Annu Rep Comp Chem 13, 231–278 (2017).

35. W. Humphrey, A. Dalke, K. Schulten, VMD: Visual molecular dynamics. Journal of Molecular Graphics & Modelling 14, 33–38 (1996).

36. Y. Miao, W. Sinko, L. Pierce, D. Bucher, J. A. McCammon, Improved reweighting of accelerated molecular dynamics simulations for free energy calculation. J Chem Theory Comput 10, 2677–2689 (2014).

