## Supporting Information for "Mechanism of Tripeptide Trimming by γ-Secretase"

**Table of Contents:**

1. **Materials and Methods (Page 3 - 12)**
2. **Supporting Figures (Figures S1 - S12) (Page 13 - 29)**
3. **Supporting Tables (Table S1) (Page 30)**
4. **References (Page 31)**

### 1. Methods and Materials

#### C100-FLAG substrates expression and purification

C100-FLAG constructs (1) were transformed into *E. coli* BL21 cells. *E. coli* BL21 cells were grown in LB media at 37$^0$C in the incubator shaker until $OD_{600}$ reached 0.6. Cells were induced with 0.5 mM IPTG and grown for 4 hours shaking at 37 ºC. The cells were then pelleted and resuspended in lysis buffer composed of 50 mM HEPES pH 8 with 1% Triton X-100. The cells were lysed by French press three times and lysate was centrifuged to remove cell debris. The clear lysate was mixed with anti-FLAG M2-agarose beads (Sigma-Aldrich) for 16 h at 4 ºC. Substrates were eluted from the beads with 100 mM glycine pH 2.5 with 0.25% NP-40, following by washing of the beads 3 times with lysis buffer. The elute was neutralized with Tris HCl and stored at -80$^0$C.

#### γ-Secretase assays

γ-Secretase purification and assays were carried out as described previously (2). Briefly, 30 nM γ-secretase was incubated for 30 min at 37 °C in assay buffer composed of 50 mM Hepes pH 7.0, 150 mM NaCl, and 0.25% 3-[(3-cholamidopropyl)dimethylammonio]-2-hydroxy-1-propanesulfonate (CHAPSO) detergent supplemented with 0.1% phosphatidylcholine (DOPC) and 0.025% phosphatidylethanolamine (DOPE). Reactions were initiated by addition purified C100-FLAG substrate to a final concentration of 5 μM and performed by incubating at 37 °C for 16 h.

#### Detection of AICD species

After 16 h, AICD-FLAG produced from the enzymatic assay was isolated by immunoprecipitation. The assay mixture was incubated with anti-FLAG M2 beads (SIGMA) in 10 mM MES pH 6.5, 10 mM NaCl, 0.05% DDM detergent for 16 h at 4 ºC. AICD products were eluted from the anti-FLAG beads with acetonitrile:water (1:1) with 0.1% trifluoroacetic acid. The elutes were run on a Bruker autoflex MALDI-TOF mass spectrometer in linear mode.

**Western blotting**

Samples from γ-secretase assays and C100-FLAG standards were run on 4-12% Bis-Tris gel and transferred to PVDF membrane. The membrane was treated with 5% dry milk in PBS Tween-20 for 1 h at ambient temperature. The membrane was then incubated with anti-FLAG M2 antibodies at 4 ºC overnight. The membrane was washed 3 times with PBS Tween-20 and incubated with anti-mouse secondary antibodies for 1 h. The membrane was washed and imaged for chemiluminescence and band signal intensity was measured by densitometry.

**LC-MS/MS tandem mass spectrometry**

Small peptides were analyzed using an ESI Quadrupole Time-of-Flight (Q-TOF) mass spectrometer (Q-TOF Premier, Waters) by LC-MS/MS experiment, as previously described (3). Briefly, assay samples and standard peptides were loaded onto a C18 column and eluted with a step gradient of 0.08% aqueous formic acid (A), acetonitrile (B), isopropanol (C), and a 1:1 acetone/dioxane mixture (D). The gradient well separated the lipids and detergent present in the buffer from the small peptides. The three most abundant collison-induced dissociation (CID) fragments were identified from the MS/MS for each small peptide. The peptide chromatographic area was obtained from the summed signals from three most abundant ions.

**Peptide Gaussian accelerated molecular dynamics (Pep-GaMD)**

Pep-GaMD is an enhanced sampling technique that works by adding a selective harmonic boost potential to smooth biomolecular potential energy surface and reduce the system energy barriers. Detail of the Pep-GaMD method has been described in previous study (4). Here, we briefly descript the algorithm of the selective Pep-GaMD method used in the current study.

We consider a system of peptide $L$ binding to a protein $P$ in a biological environment $E$. The system comprises of $N$ atoms with their coordinates $r \equiv \{\vec{r}_1, \cdots, \vec{r}_N\}$ and momenta $p \equiv \{\vec{p}_1, \cdots, \vec{p}_N\}$. The system Hamiltonian can be expressed as:

$$H(r,p) = K(p) + V(r), \quad (1)$$

where $K(p)$ and $V(r)$ are the system kinetic and total potential energies, respectively. Next, we decompose the potential energy into the following terms:

$$\begin{aligned} V(r) &= V_{P,b}(r_P) + V_{L,b}(r_L) + V_{E,b}(r_E) \\ &+ V_{PP,nb}(r_P) + V_{LL,nb}(r_L) + V_{EE,nb}(r_E) \\ &+ V_{PL,nb}(r_{PL}) + V_{PE,nb}(r_{PE}) + V_{LE,nb}(r_{LE}), \end{aligned} \quad (2)$$

where $V_{P,b}$, $V_{L,b}$ and $V_{E,b}$ are the bonded potential energies in protein $P$, peptide $L$ and environment $E$, respectively. $V_{PP,nb}$, $V_{LL,nb}$ and $V_{EE,nb}$ are the self non-bonded potential energies in protein $P$, peptide $L$ and environment $E$, respectively. $V_{PL,nb}$, $V_{PE,nb}$ and $V_{LE,nb}$ are the non-bonded interaction energies between $P$-$L$, $P$-$E$ and $L$-$E$, respectively. According to classical molecular mechanics force fields (5, 6), the non-bonded potential energies are usually calculated as:

$$V_{nb} = V_{elec} + V_{vdW}, \tag{3}$$

where $V_{elec}$ and $V_{vdW}$ are the system electrostatic and van der Waals potential energies.

The bonded potential energies are usually calculated as

$$V_b = V_{bond} + V_{angle} + V_{dihedral} \quad (4)$$

where $V_{bond}$, $V_{angle}$ and $V_{dihedral}$ are the system bond, angle and dihedral potential energies. Presumably, peptide binding mainly involves in both the bonded and non-bonded interaction energies of the peptide since peptides often undergo large conformational changes during binding to the target proteins. Among the bonded potential, the dihedral potential energy ($V_{dihedral}$) plays a critical important role in system conformational change. Thus, the essential peptide potential energy in the current study is $V_L(r) = V_{LL,\mathrm{dihedral}}(r_L) + V_{LL,nb}(r_L) + V_{PL,nb}(r_{PL}) + V_{LE,nb}(r_{LE})$. In this selective Pep-GaMD, we add boost potential selectively to the essential peptide potential energy according to the GaMD algorithm:

$$\Delta V_L(r) = \begin{cases} \frac{1}{2}k_L\big(E_L - V_L(r)\big)^2, & V_L(r) < E_L \\ 0, & V_L(r) \geq E_L, \end{cases} \tag{5}$$

where $E_L$ is the threshold energy for applying boost potential and $k_L$ is the harmonic constant. The Pep-GaMD simulation parameters are derived similarly as in the previous GaMD. When $E$ is set to the lower bound as the system maximum potential energy ($E=V_{max}$), the effective harmonic force constant $k_0$ can be calculated as:

$$k_0 = \min(1.0, k_0') = \min(1.0, \frac{\sigma_0}{\sigma_V}\frac{V_{max}-V_{min}}{V_{max}-V_{avg}}), \tag{6}$$

where $V_{max}$, $V_{min}$, $V_{avg}$ and $\sigma_V$ are the maximum, minimum, average and standard deviation of the boosted system potential energy, and $\sigma_0$ is the user-specified upper limit of the standard deviation of $\Delta V$ (e.g., 10$k_B$T) for proper reweighting. The harmonic constant is calculated as $k = k_0 \cdot \frac{1}{V_{max}-V_{min}}$ with $0 < k_0 \leq 1$ . Alternatively, when the threshold energy *E* is set to its upper bound $E = V_{min} + \frac{1}{k}$, $k_0$ is set to:

$$k_0 = k_0^{"} \equiv (1 - \frac{\sigma_0}{\sigma_V}) \frac{V_{max}-V_{min}}{V_{avg}-V_{min}}, \tag{7}$$

if $k_0^{"}$ is found to be between *0* and *1*. Otherwise, $k_0$ is calculated using Eqn. (6).

In addition to selectively boosting the peptide, another boost potential is applied on the protein and solvent to enhance conformational sampling of the protein and facilitate peptide binding. The second boost potential is calculated using the total system potential energy other than the essential peptide potential energy as:

$$\Delta V_D(r) = \begin{cases} \frac{1}{2} k_D \left(E_D - V_D(r)\right)^2, & V_D(r) < E_D \\ 0, & V_D(r) \geq E_D \end{cases} \tag{8}$$

Where $V_D$ is the total system potential energy other than the essential peptide potential energy, $E_D$ is the corresponding threshold energy for applying the second boost potential and $k_D$ is the harmonic constant. This leads to dual-boost Pep-GaMD with the total boost potential $\Delta V(r) = \Delta V_L(r) + \Delta V_D(r)$.

**Energetic Reweighting of Pep-GaMD**

To calculate potential of mean force (PMF)(7) from Pep-GaMD simulations, the probability distribution along a reaction coordinate is written as $p^*(A)$. Given the boost potential $\Delta V(\vec{r})$ of each frame, $p^*(A)$ can be reweighted to recover the canonical ensemble distribution, $p(A)$, as:

$$p(A_j) = p^*(A_j)\frac{\langle e^{\beta\Delta V(\vec{r})}\rangle_j}{\sum_{i=1}^{M}\langle p^*(A_i)e^{\beta\Delta V(\vec{r})}\rangle_i},\ j = 1, \ldots, M, \tag{9}$$

where $M$ is the number of bins, $\beta = k_B T$ and $\langle e^{\beta\Delta V(\vec{r})}\rangle_j$ is the ensemble-averaged Boltzmann factor of $\Delta V(\vec{r})$ for simulation frames found in the $j^{\text{th}}$ bin. The ensemble-averaged reweighting factor can be approximated using cumulant expansion:

$$\langle e^{\beta\Delta V(\vec{r})}\rangle = exp\left\{\sum_{k=1}^{\infty}\frac{\beta^k}{k!}C_k\right\}, \tag{10}$$

where the first two cumulants are given by

$$\begin{aligned} C_1 &= \langle \Delta V \rangle, \\ C_2 &= \langle \Delta V^2 \rangle - \langle \Delta V \rangle^2 = \sigma_v^2. \end{aligned} \tag{11}$$

The boost potential obtained from Pep-GaMD simulations usually follows near-Gaussian distribution. Cumulant expansion to the second order thus provides a good approximation for computing the reweighting factor (8, 9). The reweighted free energy $F(A) = -k_B T \ln p(A)$ is calculated as:

$$F(A) = F^*(A) - \sum_{k=1}^{2}\frac{\beta^k}{k!}C_k + F_c, \tag{12}$$

where $F^*(A) = -k_B T \ln p^*(A)$ is the modified free energy obtained from Pep-GaMD simulation and $F_c$ is a constant.

**System Setup**

Pep-GaMD simulations was performed on the γ-secretase activation for ζ cleavage of Aβ49. Active APP-bound γ-secretase was taken from the previous study (2) and the amide bond between Aβ49 and AICD50-99 was cleaved as the starting structure. The enzyme is based on previously published cryo-EM structure (10) **(Fig. S1)** with Asp385 computationally restored, artificial enzyme-substrate disulfide bond removed and missing residues on APP N-terminus added. The Ala385 residue in the cryo-EM structure was computationally mutated back to Asp385. Two artificial disulfide bonds between Cys112 of PS1-Q112C and Cys4 of PS1-V24C were removed as the wildtype residues were restored. SWISS-MODEL (11) homology modeling was used to restore 5 N-terminal APP residues that were missing in the cryo-EM structure. The N- and C-termini of the receptor were capped with the acetyl (ACE) and N-methyl amide (NME) neutral groups, respectively. Protein residues were set to the standard CHARMM protonation states at neutral pH with the *psfgen* plugin in VMD (12). Then the complex was embedded in a 1-palmitoyl-2-oleoyl-sn-glycero-3-phosphocholine (POPC) bilayer with all overlapping lipid molecules removed using the *Membrane* plugin in VMD (12) **(Figure S1)**. The system charges were then neutralized at 0.15 M NaCl using the *Solvate* plugin in VMD.(12) Periodic boundary conditions were applied on the simulation systems. The simulation systems of γ-secretase bound by wildtype and mutant APP are summarized in **Table 1**. For APP-mutant simulations systems, isoleucine, alanine, valine, isoleucine and threonine residues were mutated to phenylalanine, threonine, phenylalanine, threonine and proline computationally at the $45^{th}$, $42^{nd}$, $46^{th}$, $45^{th}$ and $48^{th}$ residue of APP substrate, respectively. These corresponded to I45F, A42T, V46F, I45T and T48P mutations as per the numbering based on C99, although the actual substrate in the model was based on C83.

In addition to the γ-secretase bound by Aβ49 and charged N-terminal AICD50-99, we tested Pep-GaMD simulations on enzyme systems bound by Aβ49 in the absence and presence of neutral N-terminal AICD50-99 **(Fig. S3)**. The neutral and charged N-terminus of the AICD50-99 was characterized by the presence of $-NH_2$ and $NH_3^+$ functional groups at the N-terminal end, respectively. Unlike the charged N-terminal AICD50-99 system, activation was not observed during the 600 ns of Pep-GaMD of either of the enzyme systems bound by Aβ49 in the absence and presence of neutral N-terminal AICD50-99 **(Figs. S5E-S5F and S8E-S8F)**. Free energy profiles were plotted for the Pep-GaMD simulations of both the enzyme systems. Two low energy conformational states were identified in the system without the AICD bound including "Inhibited-2" and "Intermediate" **(Fig. S9A)**. Similarly, "Initial" and "Intermediate" low energy conformational states were identified in the free energy profile of the enzyme system bound to Aβ49 in the presence of neutral N-terminal AICD50-99 **(Fig. S9B)**. The "Inhibited-2" and the "Intermediate" conformational states here were same as the one identified in the wildtype mutant and the V46F γ-secretase system, respectively **(Fig. 2A and 2D)**. The "Initial" conformational state resembled the one identified in the wildtype, I45F and A42T mutant systems **(Fig. 2A-2C)**. In comparison, "Final" active conformational state was identified in the wildtype system bound to Aβ49 and charged N-terminal AICD50-99 **(Fig. 2A)**. Therefore, systems for γ-secretase bound by Aβ49 and charged N-terminal AICD50-99 were used for final Pep-GaMD simulations.

**Simulation Protocol**

The CHARMM36m (13) parameter set was used for the protein and POPC lipids. Initial energy minimization and thermalization of the γ-secretase complex followed the same protocol as used in the previous GaMD simulations of membrane proteins (14, 15). The simulation proceeded with equilibration of lipid tails. With all the other atoms fixed, the lipid tails were energy minimized for 1000 steps using the conjugate gradient algorithm and melted with constant number, volume, and temperature (NVT) run for 0.5 ns at 310 K. Each system was further equilibrated using constant number, pressure, and temperature (NPT) run at 1 atm and 310 K for 10 ns with 5 kcal (mol Å$^2$)$^{-1}$

harmonic position restraints applied to the protein. Further equilibration of the systems was performed using an NPT run at 1 atm and 310 K for 0.5 ns with all atoms unrestrained. Conventional MD simulation was performed on each system for 10 ns at 1 atm pressure and 310 K with a constant ratio constraint applied on the lipid bilayer in the X-Y plane. The Pep-GaMD simulations were carried out using AMBER 20 (13). Dual-boost Pep-GaMD simulations were performed to study the γ-secretase enzyme activation for ζ cleavage (**Table 1**). In the Pep-GaMD simulations, the threshold energy $E$ for adding boost potential was set to the upper bound, i.e. $E = V_{min} + (1/k)$(9, 16). The simulations included 50 ns equilibration after adding the boost potential and then multiple independent production runs lasting 600 ns with randomized initial atomic velocities. Pep-GaMD production simulation frames were saved every 0.2 ps for analysis.

**Simulation analysis**

The VMD(12) and CPPTRAJ (17) tools were used to visualize and analyze the Pep-GaMD trajectories. The distance between the catalytic aspartates was calculated between the Cγ atoms. Hydrogen bond distance was calculated between donor protonated oxygen atom of PS1 Asp257 and the acceptor carbonyl oxygen atom of APP substrate residue Val46. Root-mean-square fluctuations (RMSFs) were calculated for the protein residues, averaged over three independent Pep-GaMD simulations and color coded for schematic representation of each complex system. The CPPTRAJ was used to calculate the protein secondary structure plots. The *PyReweighting* toolkit(8) was applied to reweight Pep-GaMD simulations for free energy calculations by combining all simulation trajectories for each system. Bin size of 1-3 Å was used for the PMF calculation of distances. The cutoff was set to 500-1000 frames in each bin for calculating the 2D PMF profiles. Protein snapshots were taken every 1 ps for structural clustering. Clustering was performed on the Pep-GaMD simulations of wildtype, I45F, A42T and V46F mutant Aβ49 bound γ-secretase based on the RMSD of PS1 using hierarchical agglomerative algorithm in CPPTRAJ (17) generating ~10 representative structural clusters for each system. The top structural cluster was identified as the representative Final active conformational states for each γ-secretase system.

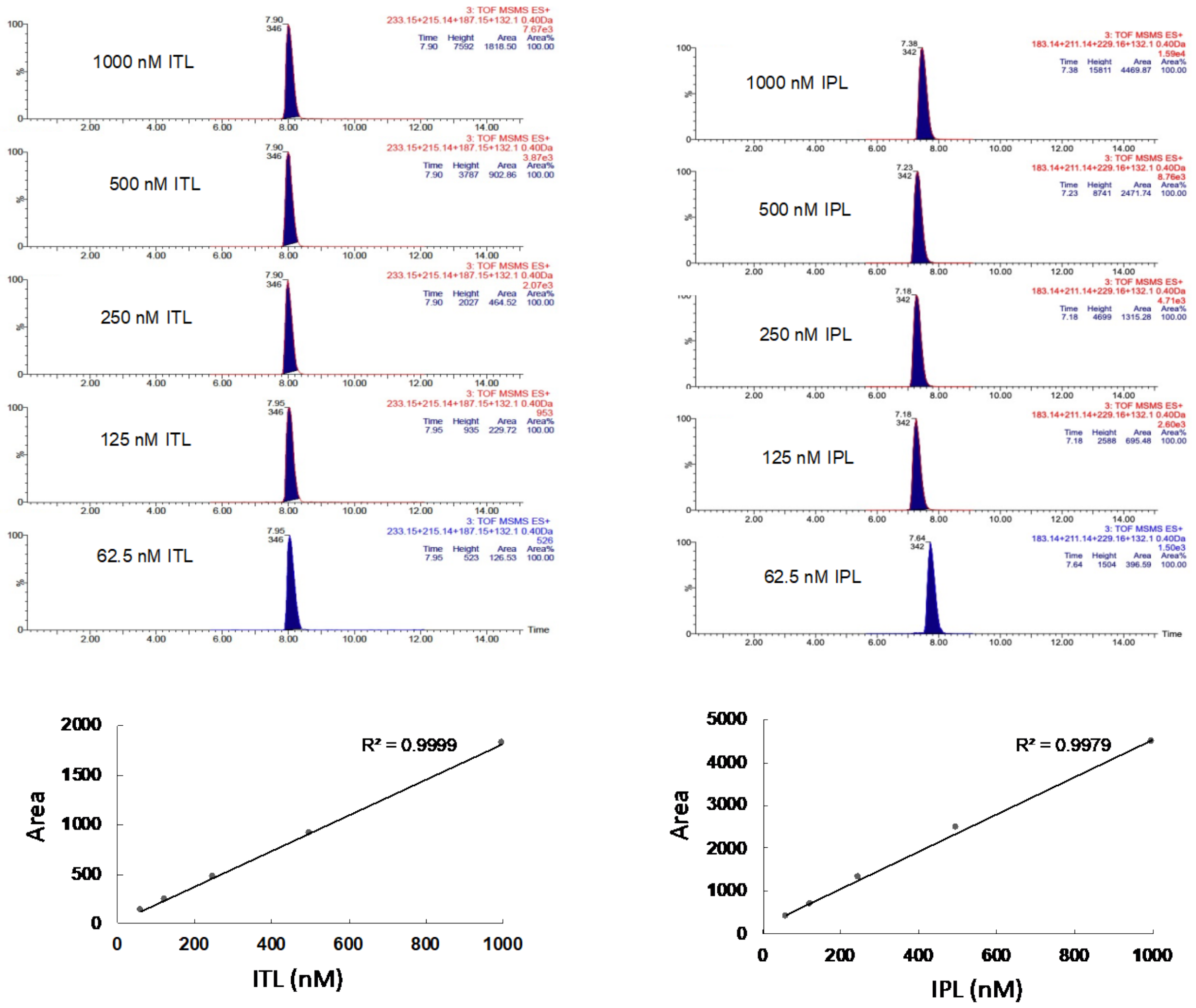


**Figure S1:** LC-MS/MS of all small-peptide standards (ITL and IPL) predicted to be generated for C100 substrates tested after γ-secretase digestion of substrates. Chromatograms are selected ion plots of the three most abundant sequence-specific product ions, selected with a 0.03 unit window. Standard curves for all small peptides were generated by plotting of the resulting peak areas of ion plots against the small-peptide concentration.

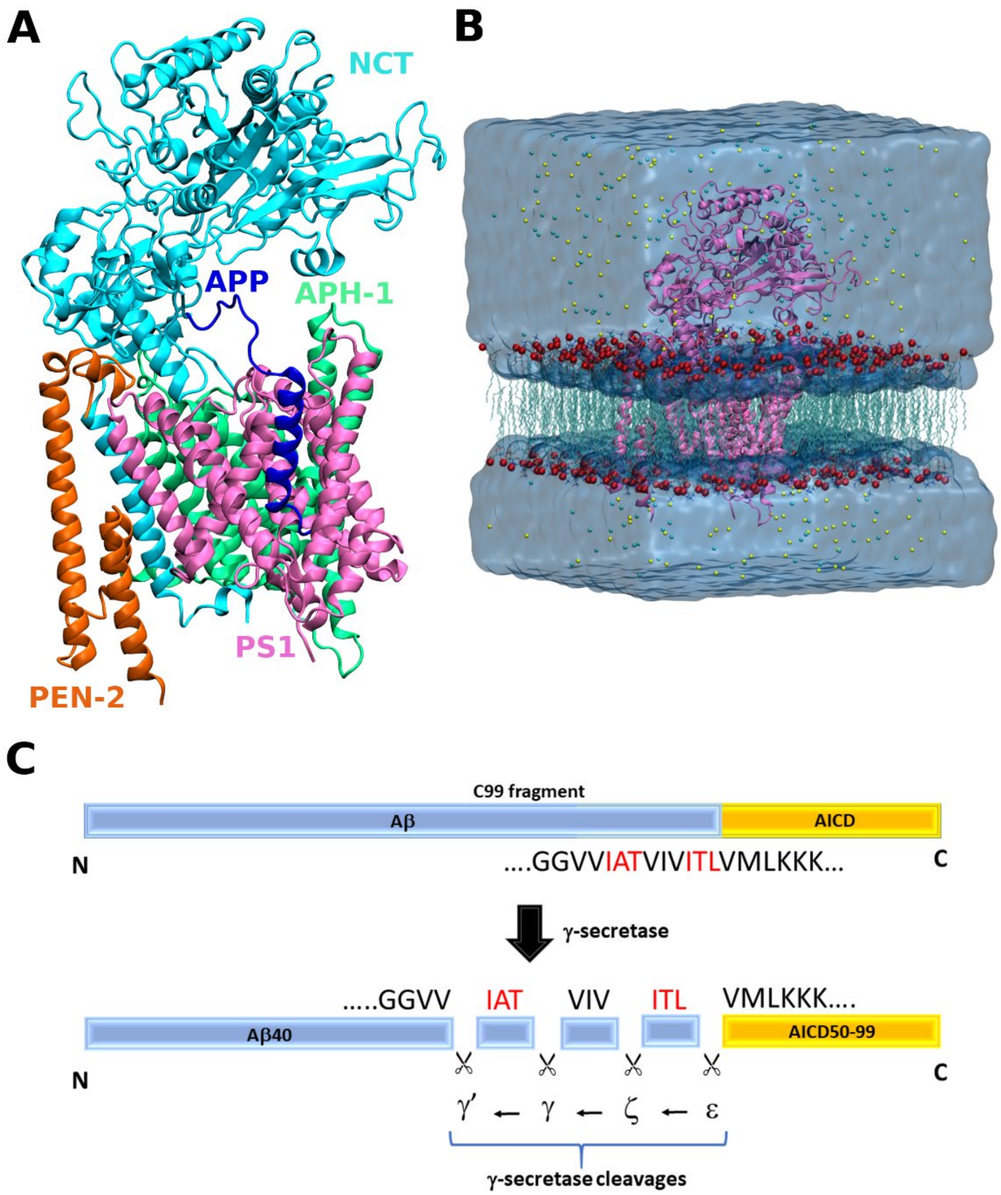


**Figure S2:** (A) γ-secretase structure bound to Aβ49 substrate (blue) with Niscastrin (NCT, cyan), Presenilin-1 (PS1, pink), Anterior Pharynx-Defective 1 (APH-1, green) and Presenilin Enhancer-2 (PEN-2, orange) subunits. The enzyme-substrate complex is represented in ribbons. (B) Pep-GaMD computational model of γ-secretase complex. The protein was embedded into a POPC lipid bilayer and solvated in an aqueous medium of 0.15 M NaCl. (C) Schematic representation of ε cleavage and processive proteolysis of APP substrate by γ-secretase.

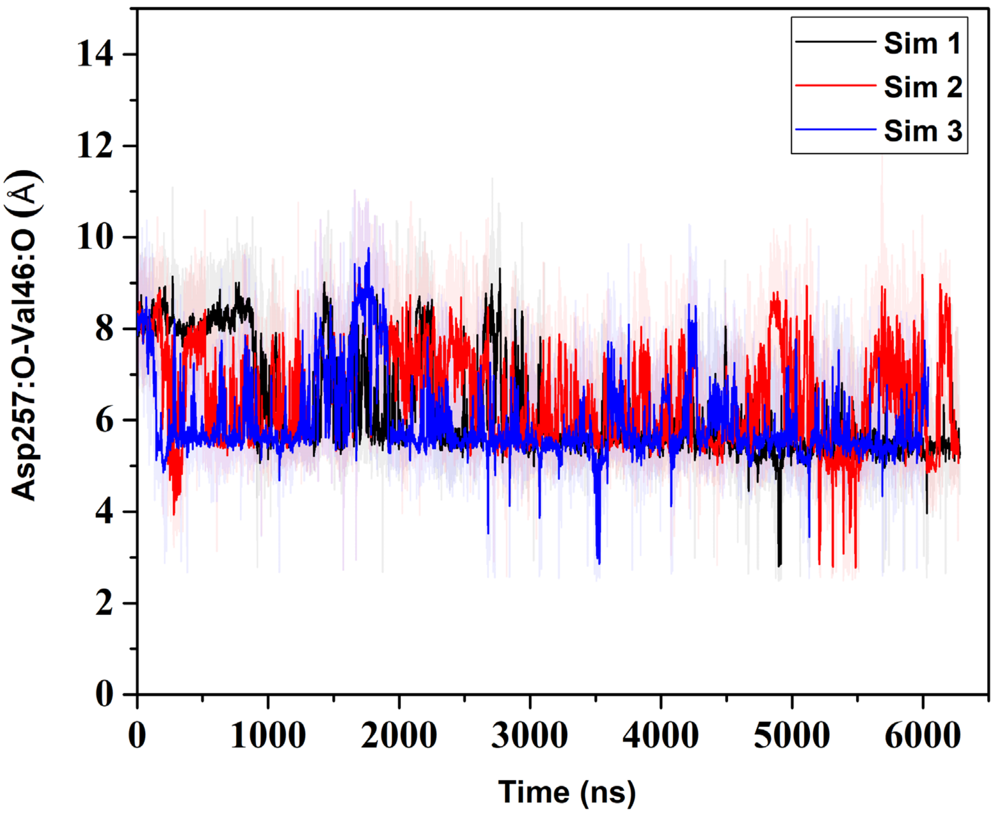


**Figure S3:** Timecourse of the Asp257:protonated O - Leu49:O distance calculated from GaMD simulations of WT γ-secretase system. More than 6 μs long GaMD simulations of the enzyme could not capture stable activation for ζ cleavage.

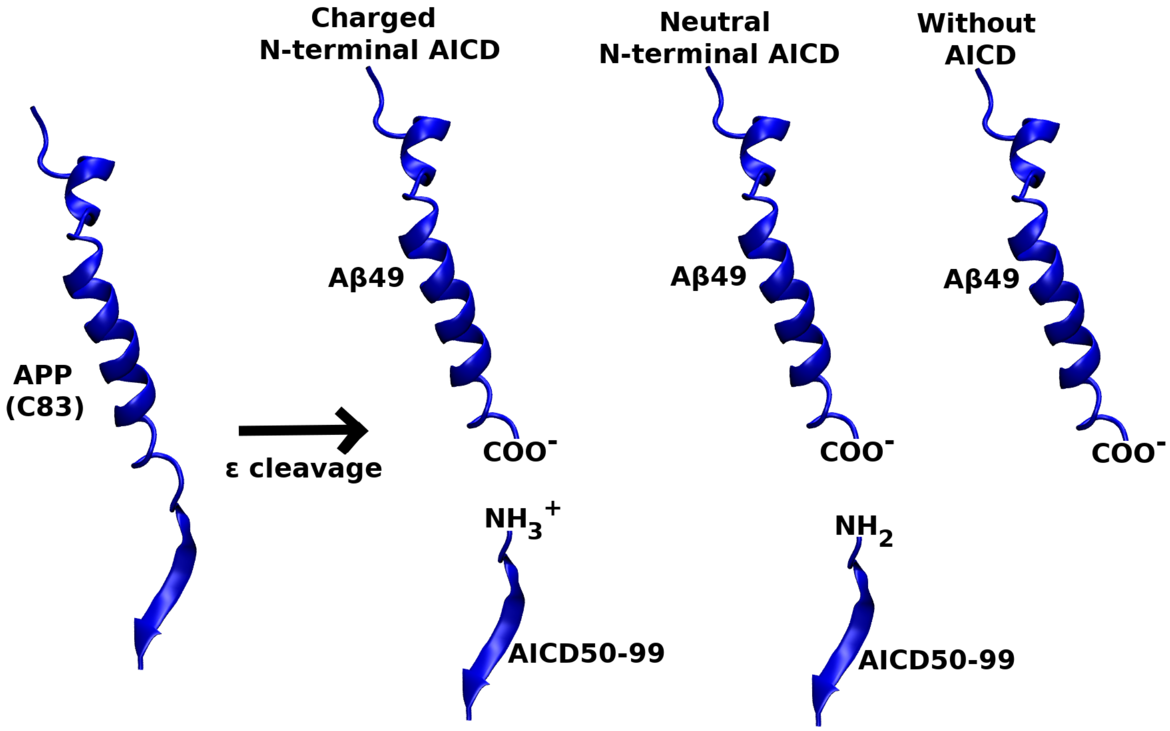


**Figure S4:** Ribbon representation of Aβ49 peptide and differently charged N-terminal AICD50-99 after ε cleavage of APP substrate by γ-secretase. Three different systems of γ-secretase bound to Aβ49 in the absence and presence of neutral and charged N-terminal AICD were used for Pep-GaMD simulations.

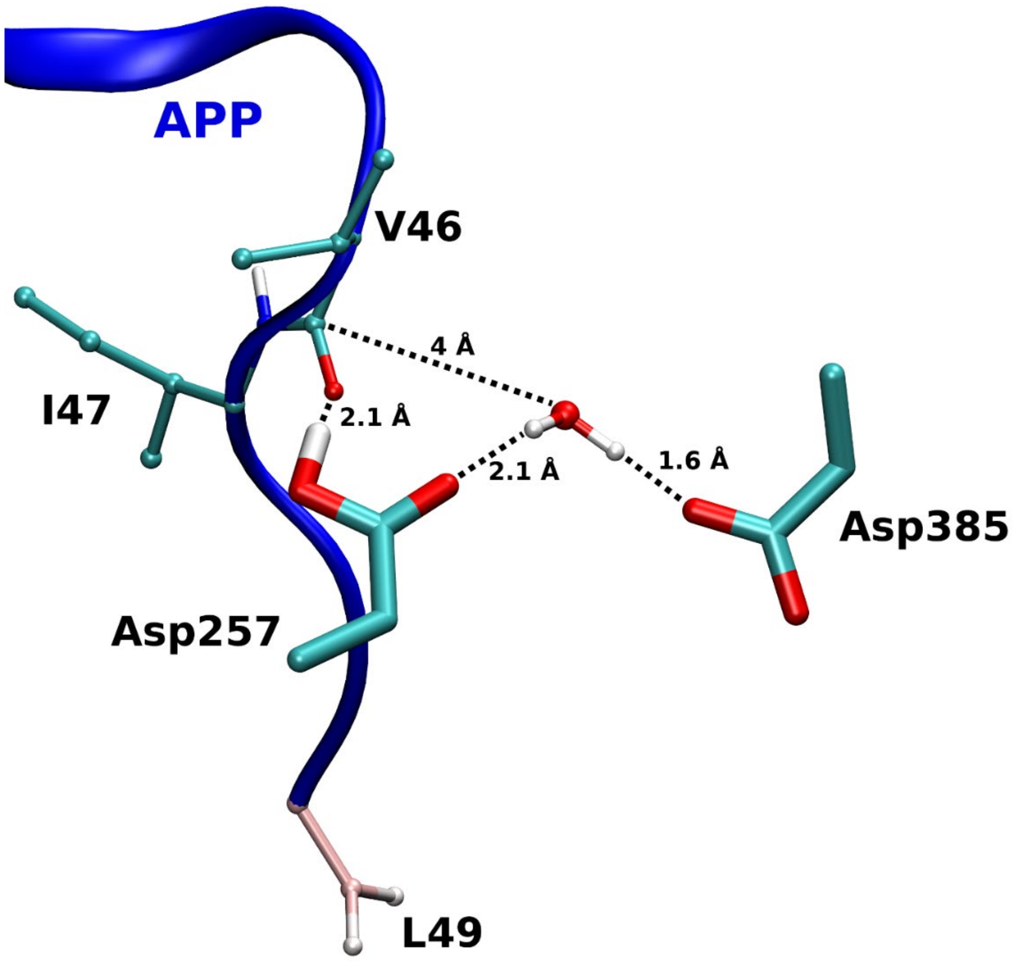


**Figure S5:** The active site poised for ζ cleavage proteolysis. The enzyme activation for ζ cleavage was characterized by coordinated hydrogen bonding between the enzyme Asp257 and carbonyl oxygen of C99 Val46. The water molecule could form hydrogen bond interactions with both catalytic aspartates and is at ~4 Å distance away from the carbonyl carbon of Val46 residue. The APP substrate (blue), aspartates and APP residues are shown as ribbon, stick and, balls and sticks, respectively.

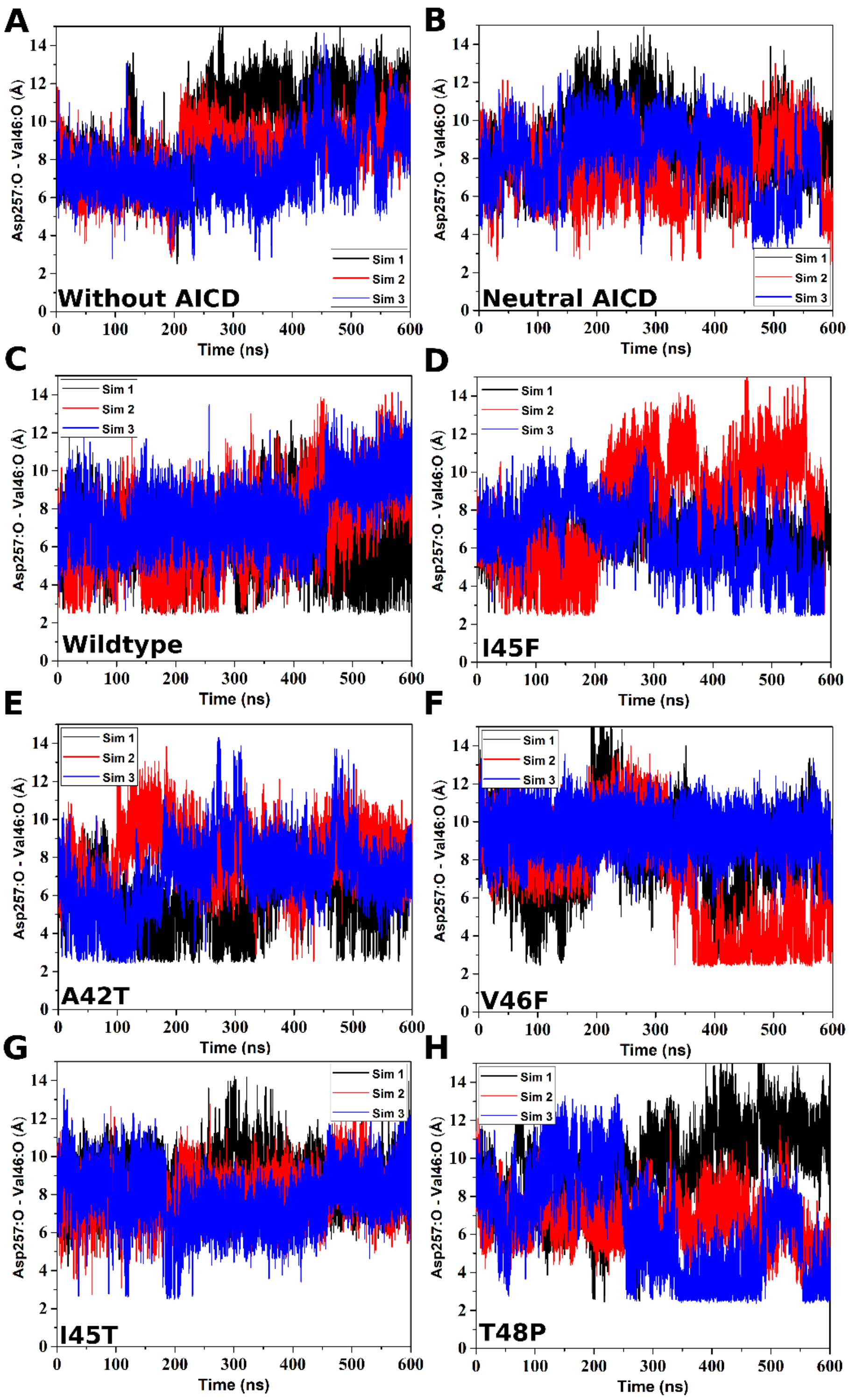


**Figure S6:** Time courses of the Asp257:protonated O - Leu49:O distances calculated from Pep-GaMD simulations of (A) WT without AICD , (B) WT with neutral AICD, (C) WT, (D) I45F, (E) A42T, (F) V46F, (G) I45T and (H) T48P APP, all with N-terminally charged AICD, bound γ-secretase systems.

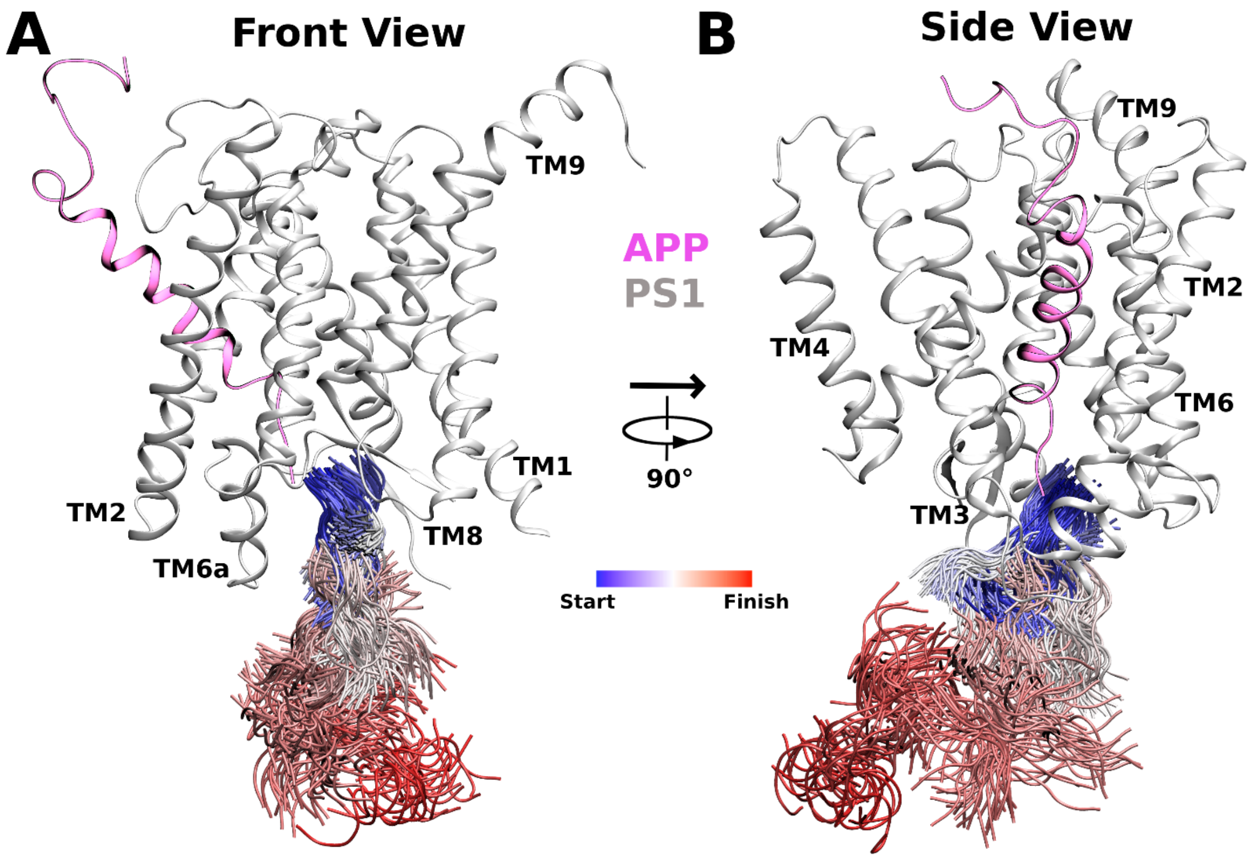


**Figure S7:** AICD50-99 dissociation pathway observed in Pep-GaMD simulations of γ-secretase system bound to wildtype APP colored by simulations time in a blue-white-red (BWR) color scheme.

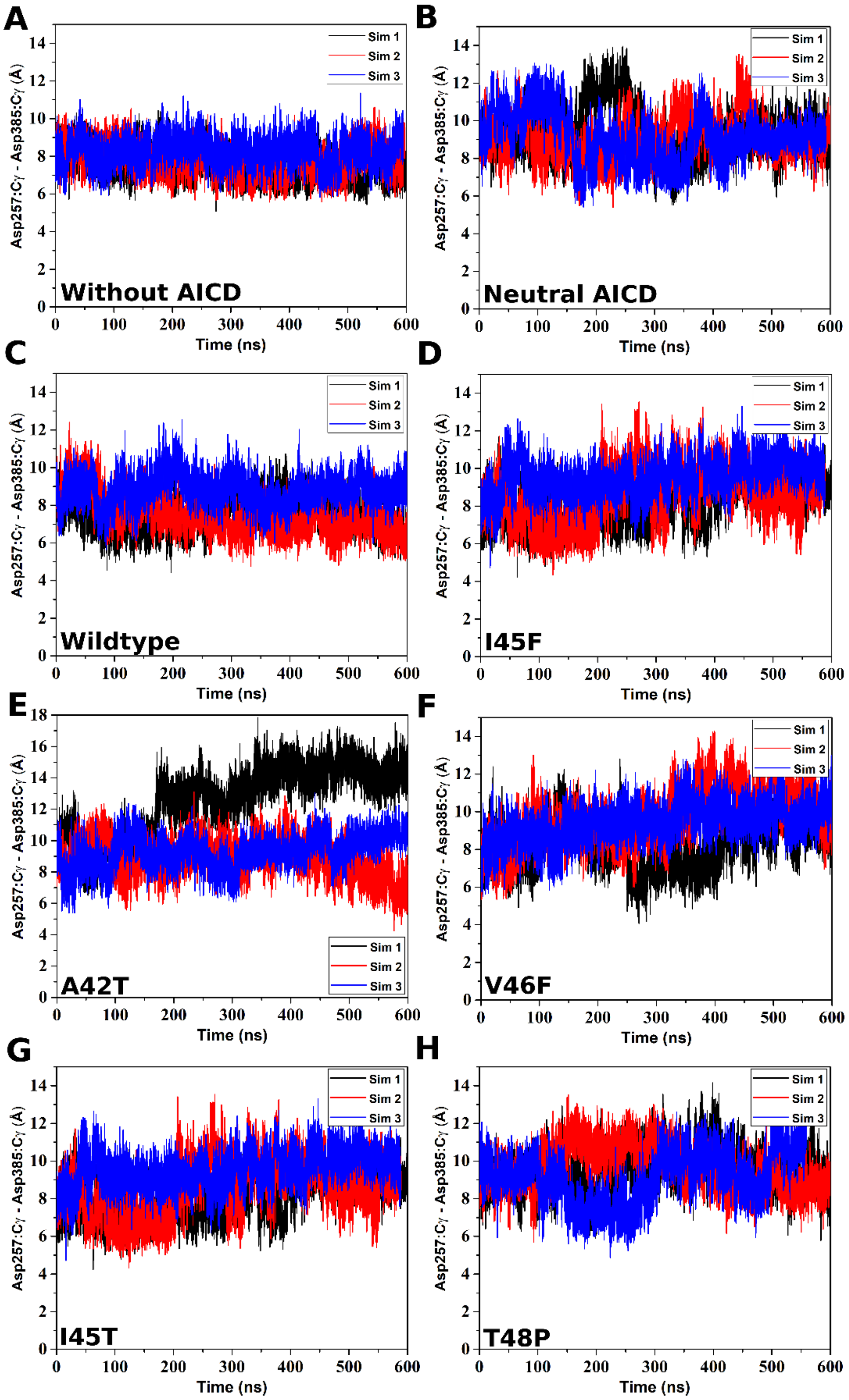


**Figure S8:** Time courses of the Asp257:Cγ - Asp385:Cγ distances calculated from Pep-GaMD simulations of (A) WT without AICD , (B) WT with neutral AICD, (C) WT, (D) I45F, (E) A42T, (F) V46F, (G) I45T and (H) T48P APP bound γ-secretase system.

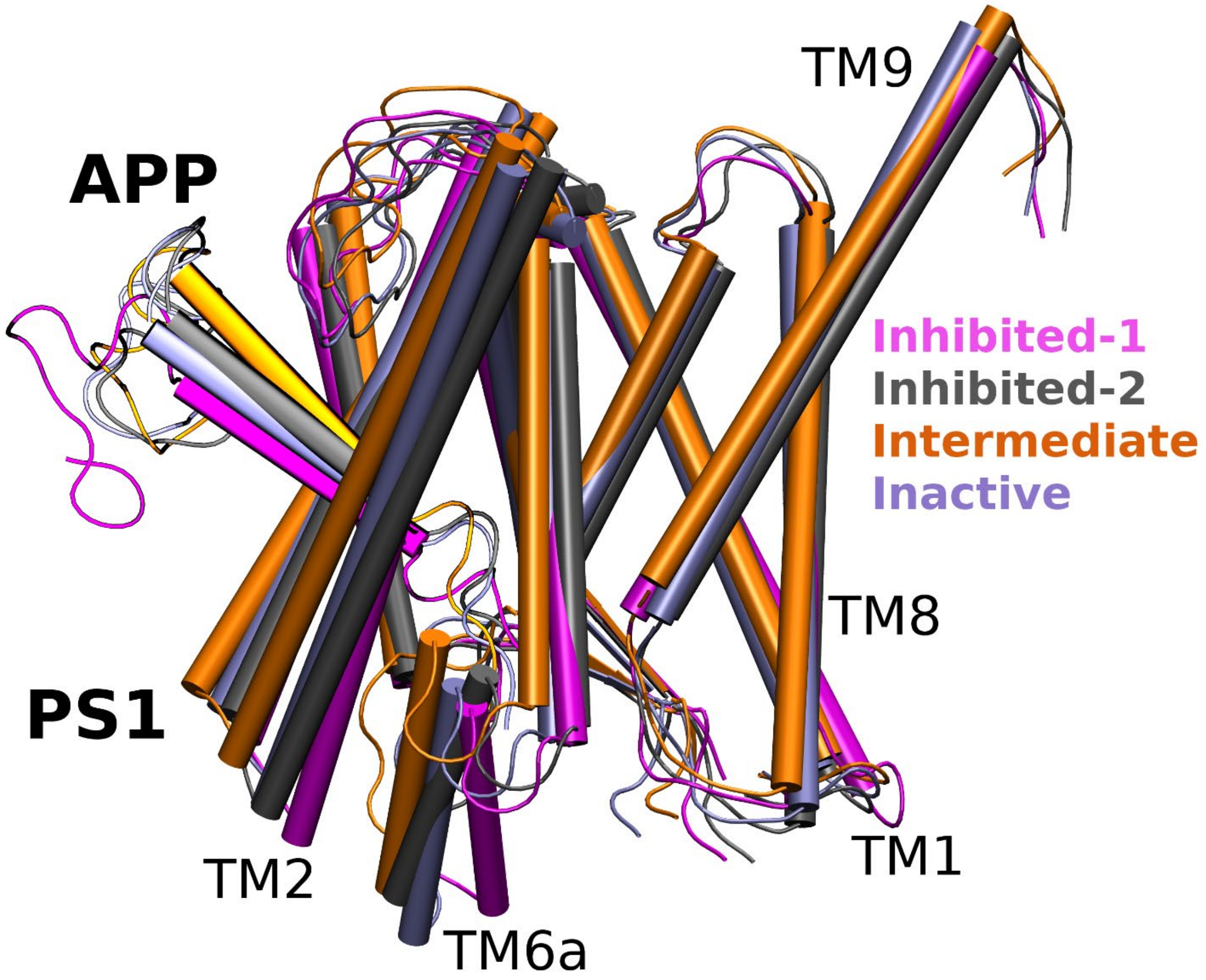


**Figure S9:** Comparison of different low energy state conformations as identified from Pep-GaMD free energy profiles of wildtype and mutant APP bound γ-secretase systems including Inhibited-1 (magenta), Inhibited-2 (gray), Intermediate (orange) and Inactive (ice blue) states.

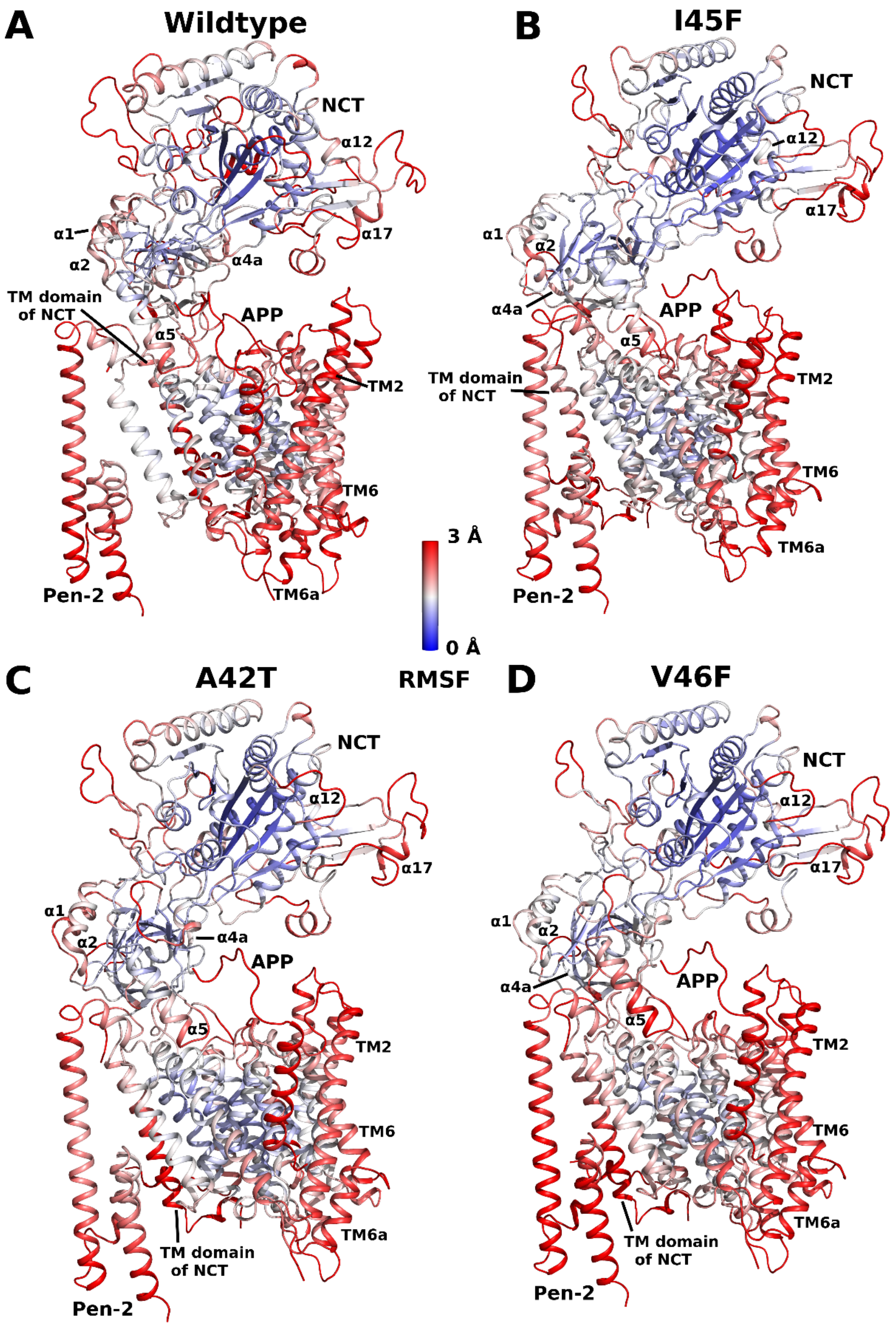


**Figure S10:** Root mean square fluctuation (RMSF) plots of different γ-secretase systems bound to (A) wildtype, (B) I45F mutant, (C) A42T mutant, and (D) V46F mutant APP. The RMSF is shown in blue-white-red color scheme for 0-3 Å of fluctuations in the enzyme-substrate complex.

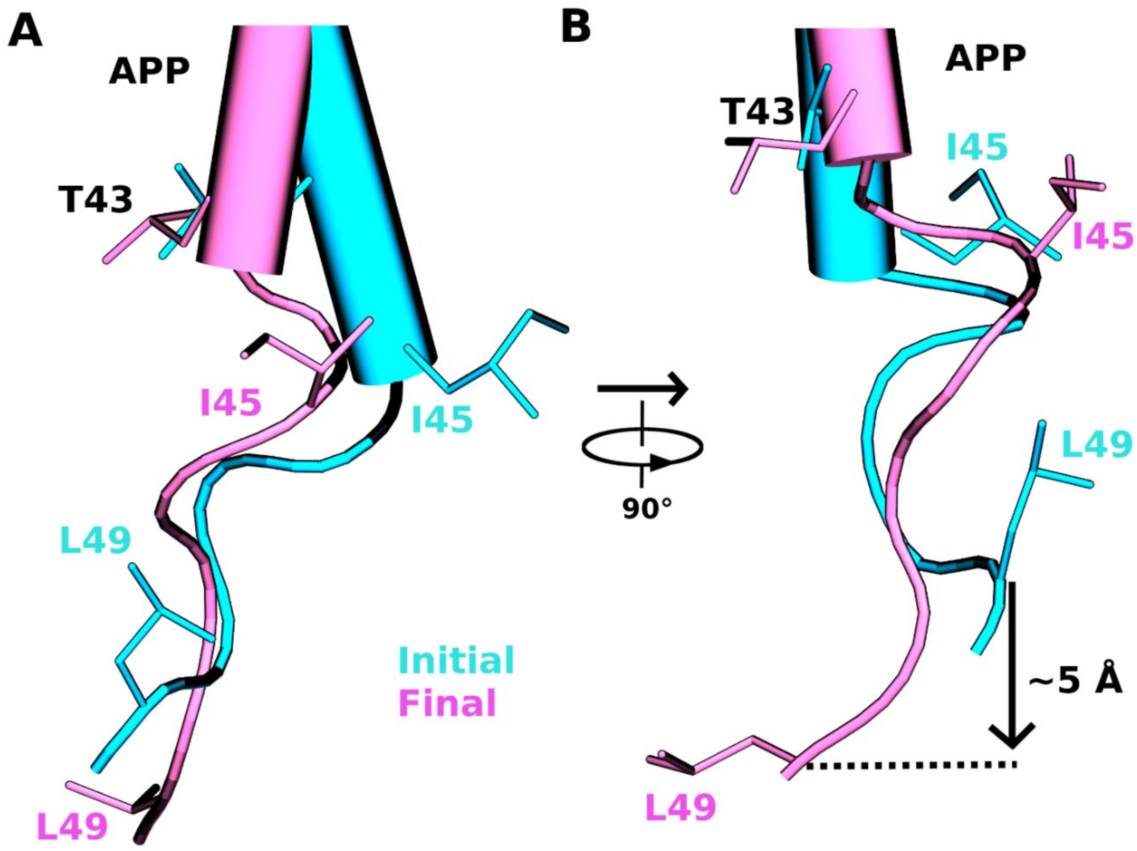


**Figure S11: (A)** Front and **(B)** side view comparison of relative positions of APP residues T43, I45 and L49 in the Initial and Final active states of the γ-secretase.

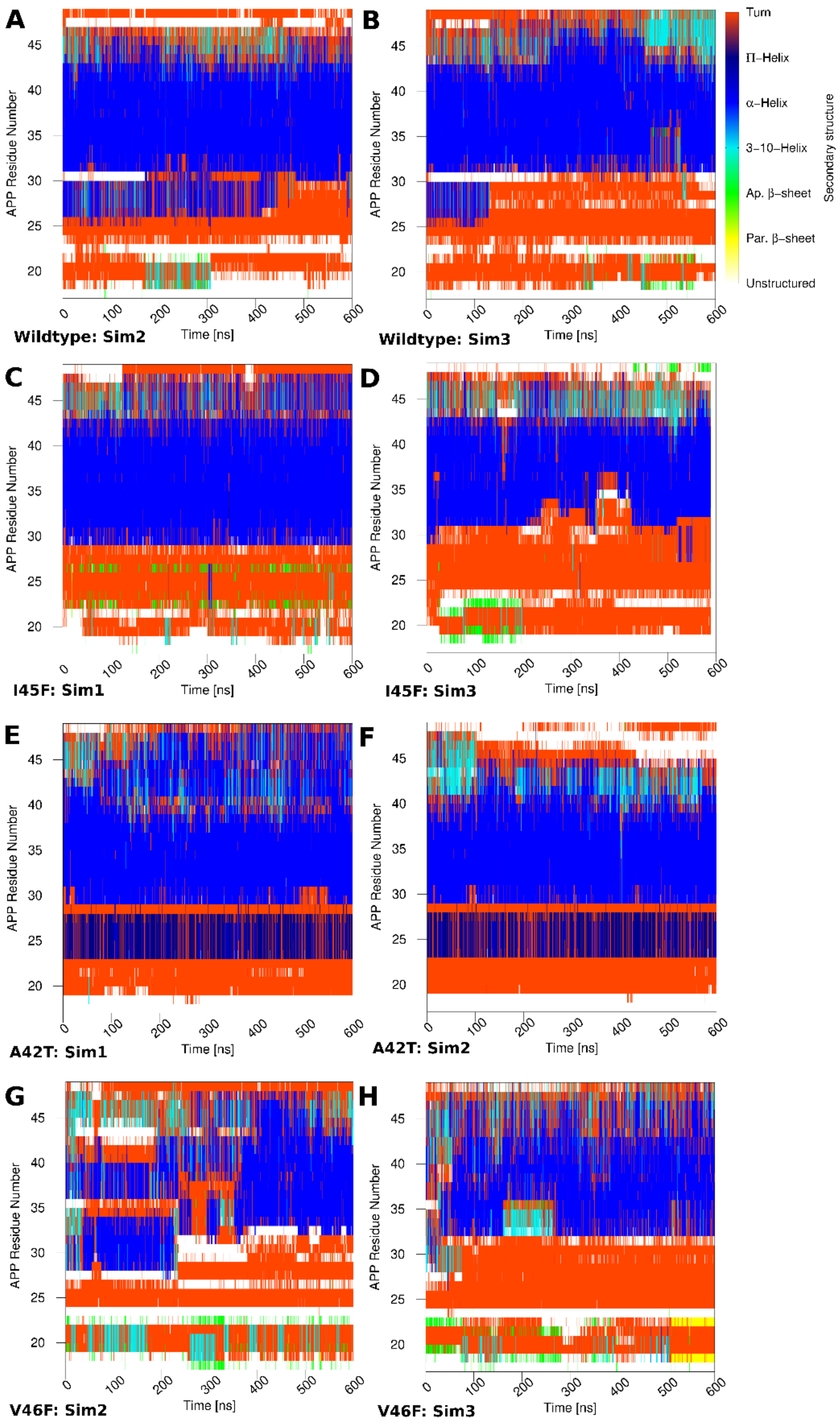


**Figure S12:** Time courses of secondary structures changes in the (A) Sim 2 and (B) Sim3 Pep-GaMD simulations of WT, (C) Sim 1 and (D) Sim 3 Pep-GaMD simulations of I45F, (E) Sim1 and (F) Sim2 Pep-GaMD simulations of A42T, (G) Sim2 and (H) Sim3 Pep-GaMD simulations of V46F mutant APP bound to γ-secretase.

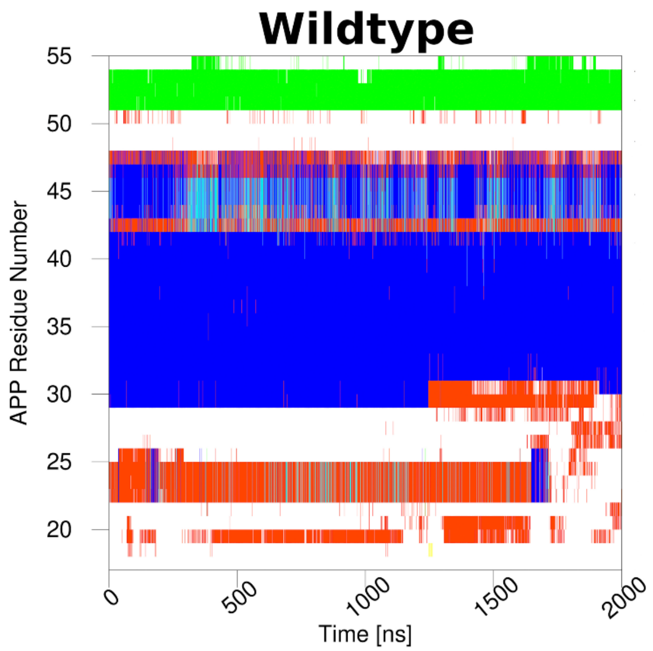


**Figure S13:** Time course of secondary structure changes in the GaMD simulations of WT APP substrate-bound γ-secretase recorded during enzyme activation for ε cleavage. This plot is extracted from our previous study(2).

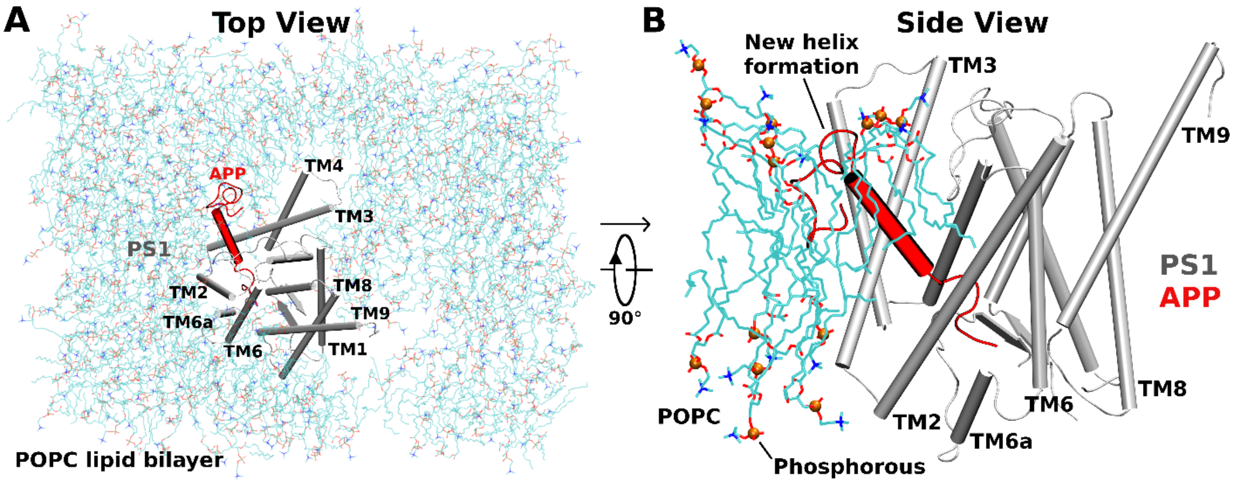


**Figure S14:** (A) Top and (B) Side view of APP (Aβ49) bound γ-secretase PS1 interacting with the POPC lipid bilayer membrane. The N-terminus of APP substrate during the ζ cleavage activation bends and interacts with the hydrophobic lipid bilayer to form α-helix conformation.

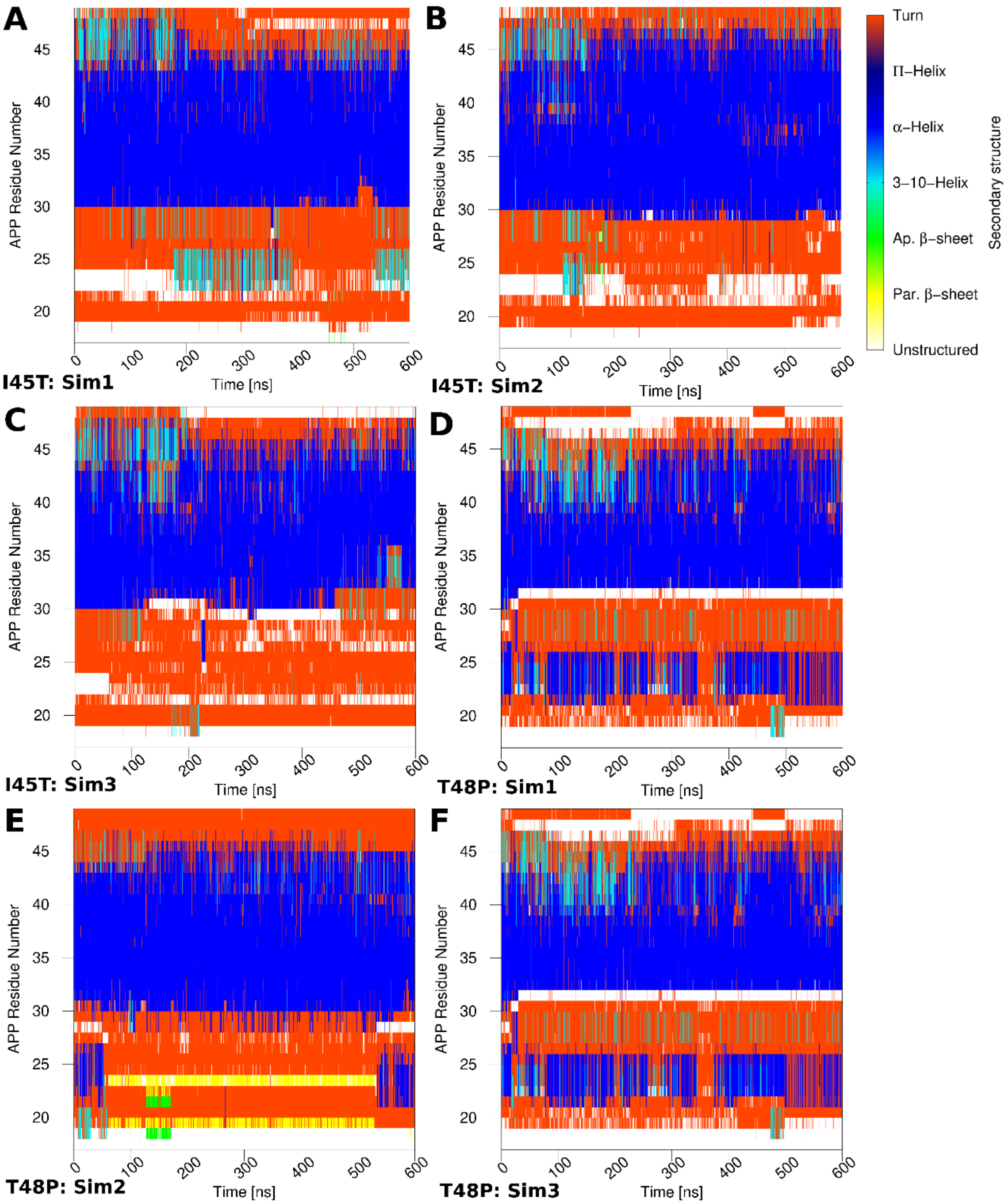


**Figure S15:** Time courses of secondary structure changes in the (A) Sim 1, (B) Sim2 and (C) Sim3 Pep-GaMD simulations of I45T mutant and (D) Sim 1, (E) Sim2 and (F) Sim3 Pep-GaMD simulations of T48P mutant APP bound to γ-secretase.

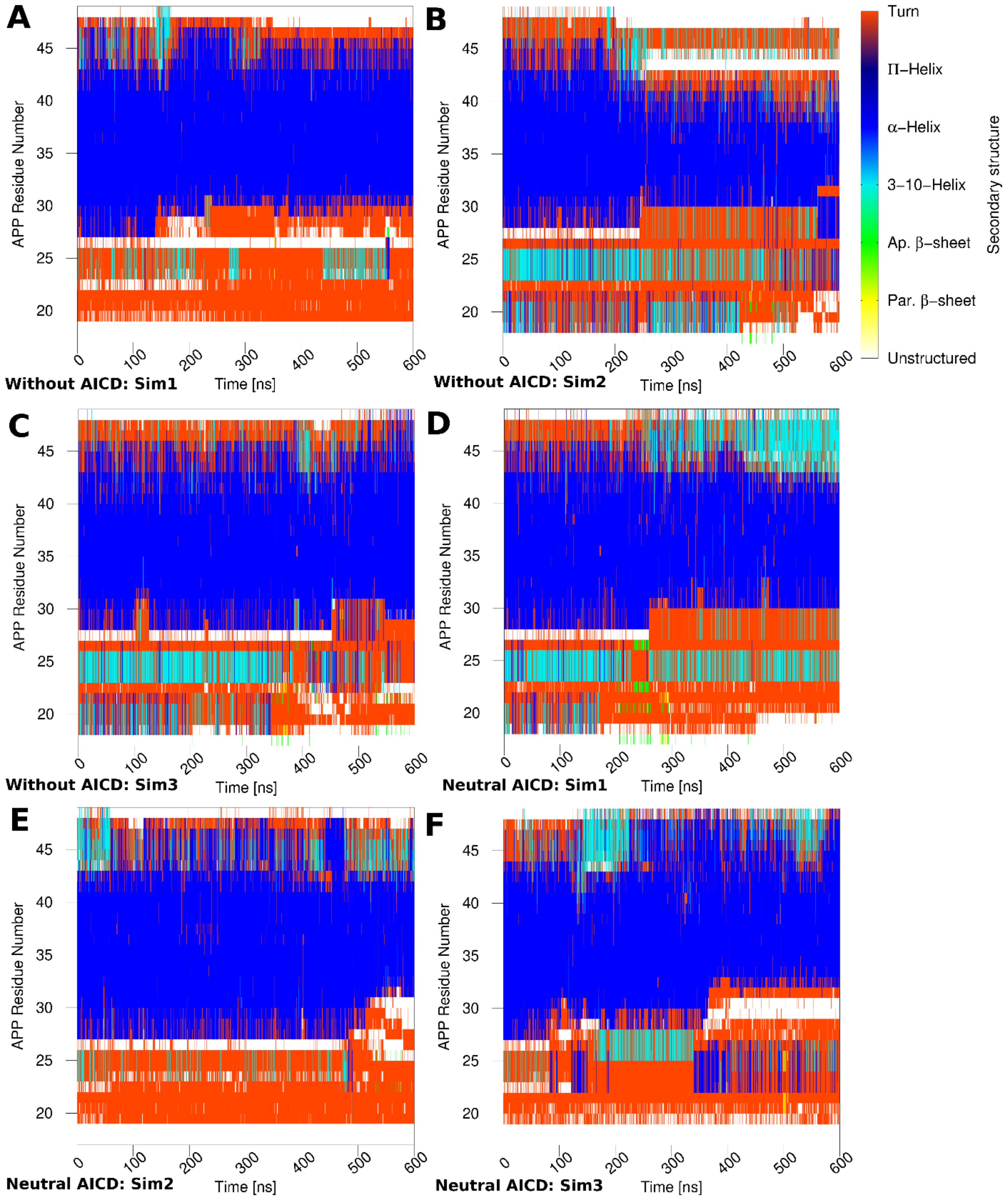


**Figure S16:** Time courses of secondary structure changes in the (A) Sim 1, (B) Sim2 and (C) Sim3 Pep-GaMD simulations of wildtype without AICD and (D) Sim 1, (E) Sim2 and (F) Sim3 Pep-GaMD simulations of wildtype with neutral AICD bound to γ-secretase.

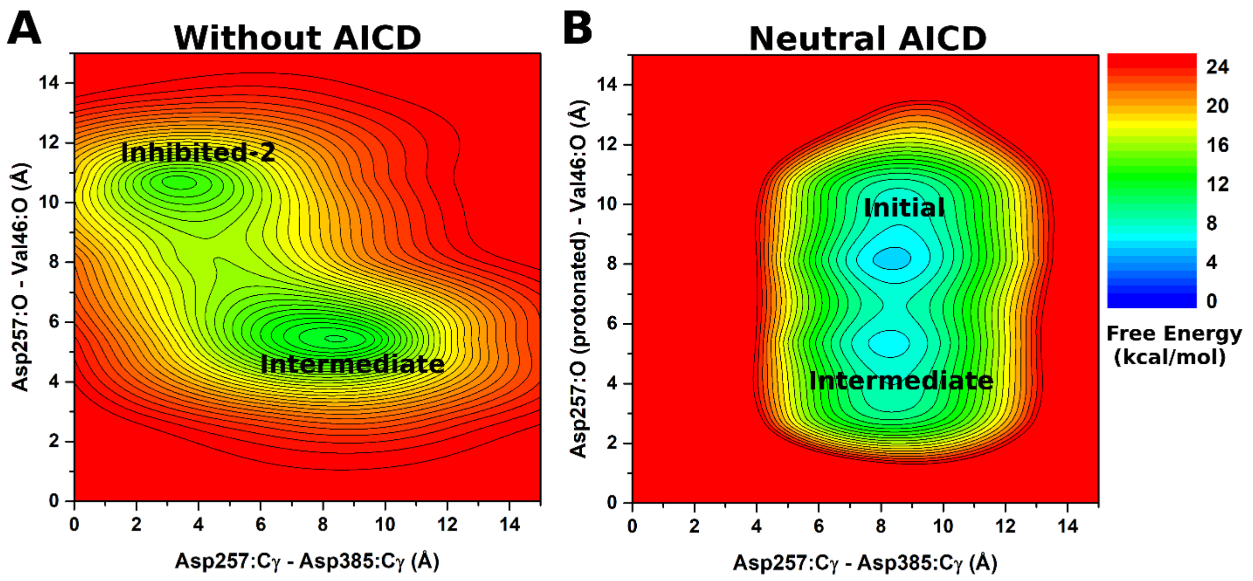


**Figure S17:** 2D free energy profiles of the Asp257:Cγ - Asp385:Cγ and Asp257:protonated O - Leu49:O distances calculated from Pep-GaMD simulations of (A) wildtype without AICD and (B) wildtype with neutral AICD bound γ-secretase system.

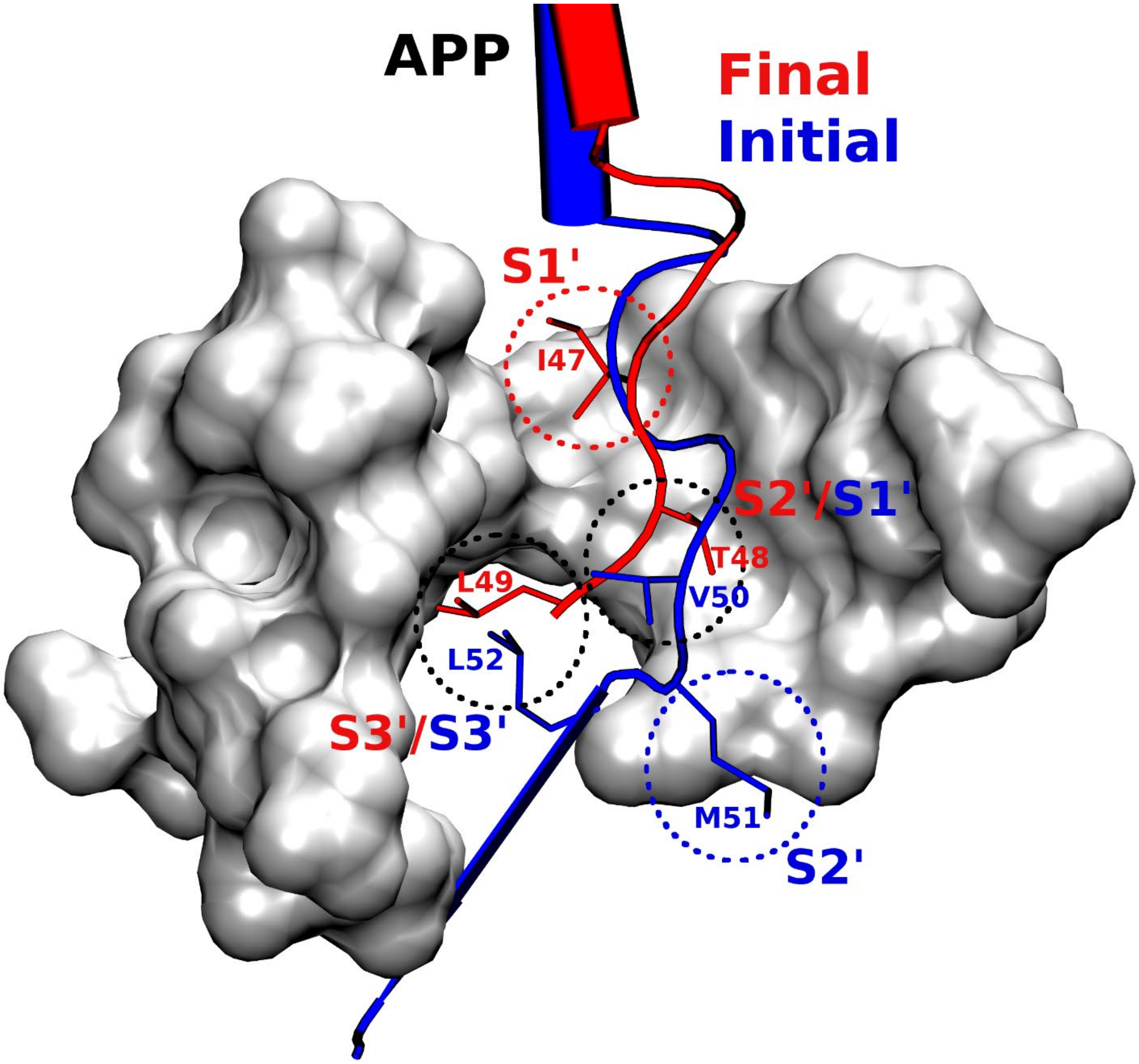


**Figure S18**: Comparison of active site subpockets of the Final active state during ζ cleavage to that of the Initial active state during ε cleavage.

**Table S1:** Residues constituting the active site subpockets S1', S2' and S3' occupied by P1', P2' and P3' Aβ49 residues for different γ-secretase systems bound to wildtype and I45F, A42T and V46F FAD mutant Aβ49.

| System | S1' | S2' | S3' |
|---|---|---|---|
| **Wildtype/I45F/A42T** | L249<br>Y256<br>L268<br>I287<br>L286<br>L271<br>L272<br>L150<br>T147<br>L282<br>F283<br>W165<br>V261<br>G382<br>L383<br>G384 | P433<br>L435<br>L258<br>V261<br>L268<br>K380<br>L381<br>G382 | L268<br>L271<br>V272<br>A275<br>L282<br>F283<br>I287<br>L381<br>G382<br>L383<br>I287 |
| **V46F** | I253<br>T147<br>Y256<br>L268<br>L271<br>M146 | L249<br>Y256<br>L268<br>I287<br>L286<br>L271<br>L272<br>L150<br>T147<br>L282<br>F283<br>W165<br>V261<br>G382<br>L383<br>G384 | L268<br>L271<br>V272<br>A275<br>L282<br>F283<br>I287<br>L381<br>G382<br>L383<br>I287 |

### Reference


1. Li Y-M*, et al.* (2000) Presenilin 1 is linked with γ-secretase activity in the detergent solubilized state. *Proceedings of the National Academy of Sciences* 97(11):6138-6143.
2. Bhattarai A, Devkota S, Bhattarai S, Wolfe MS, & Miao Y (2020) Mechanisms of γ-Secretase Activation and Substrate Processing. *ACS Central Science*.
3. Devkota S, Williams TD, & Wolfe MS (2021) Familial Alzheimer's disease mutations in amyloid protein precursor alter proteolysis by γ-secretase to increase amyloid β-peptides of≥ 45 residues. *Journal of Biological Chemistry* 296:100281.
4. Wang J & Miao Y (2020) Peptide Gaussian accelerated molecular dynamics (Pep-GaMD): Enhanced sampling and free energy and kinetics calculations of peptide binding. *J Chem Phys* 153(15):154109.
5. Vanommeslaeghe K & MacKerell AD, Jr. (2015) CHARMM additive and polarizable force fields for biophysics and computer-aided drug design. *Biochim. Biophys. Acta.* 1850(5):861-871.
6. Duan Y*, et al.* (2003) A point-charge force field for molecular mechanics simulations of proteins based on condensed-phase quantum mechanical calculations. *J. Comput. Chem.* 24(16):1999-2012.
7. Roux B (1995) The Calculation of the Potential of Mean Force Using Computer-Simulations. *Comput Phys Commun* 91(1-3):275-282.
8. Miao Y, Sinko W, Pierce L, Bucher D, & McCammon JA (2014) Improved reweighting of accelerated molecular dynamics simulations for free energy calculation. *J Chem Theory Comput* 10(7):2677–2689.
9. Miao Y, Feher VA, & McCammon JA (2015) Gaussian Accelerated Molecular Dynamics: Unconstrained Enhanced Sampling and Free Energy Calculation. *J Chem Theory Comput* 11(8):3584-3595.
10. Zhou R*, et al.* (2019) Recognition of the amyloid precursor protein by human γ-secretase. *Science* 363(6428).
11. Waterhouse A*, et al.* (2018) SWISS-MODEL: homology modelling of protein structures and complexes. *Nucleic acids research* 46(W1):W296-W303.
12. Humphrey W, Dalke A, & Schulten K (1996) VMD: Visual molecular dynamics. *Journal of Molecular Graphics & Modelling* 14(1):33-38.
13. Case DA*, et al.* (2020) Amber 2020.
14. Kappel K, Miao Y, & McCammon JA (2015) Accelerated Molecular Dynamics Simulations of Ligand Binding to a Muscarinic G-protein Coupled Receptor. *Quarterly Reviews of Biophysics* 48(04):479-487.
15. Miao Y & McCammon JA (2017) Gaussian Accelerated Molecular Dynamics: Theory, Implementation and Applications. *Annu Rep Comp Chem* 13:231-278.
16. Pang YT, Miao Y, Wang Y, & McCammon JA (2017) Gaussian Accelerated Molecular Dynamics in NAMD. *J Chem Theory Comput* 13(1):9-19.
17. Roe DR & Cheatham TE (2013) PTRAJ and CPPTRAJ: Software for Processing and Analysis of Molecular Dynamics Trajectory Data. *J Chem Theory Comput* 9(7):3084-3095.
